# The histone demethylase KDM3 prevents auto-immune piRNAs production in *Drosophila*

**DOI:** 10.1101/2022.06.02.494511

**Authors:** Karine Casier, Julie Autaa, Nathalie Gueguen, Valérie Delmarre, Marie P. Pauline, Stéphane Ronsseray, Clément Carré, Emilie Brasset, Laure Teysset, Antoine Boivin

## Abstract

In animals, genome integrity of the germ line is protected from transposable element (TE) activity by small, non-coding, dedicated RNAs acting as an immune system against TEs, and called PIWI-interacting RNAs (piRNAs) ^1,2^. In Drosophila, the production of piRNAs is initiated from heterochromatic loci containing remnants of TEs and enriched in histone H3 trimethylated on lysine 9 (H3K9me3) ^3–5^. These loci, called piRNA clusters, constitute a memory of past TE invasions. Little is known about how piRNA clusters are genetically defined. Using a genetic screen combined with a bimodal epigenetic state piRNA cluster (*BX2*), we identified the splicing factor Half pint (Hfp) and the histone demethylase KDM3 as being able to prevent *BX2* piRNA production. Furthermore, we showed that Hfp is needed to splice *Kdm3* transcripts. Germline expression of *Kdm3* coding sequence (splicing-independent) rescued the *hfp* germline knock-down (GLKD) effect demonstrating that *Kdm3* is sufficient to prevent *BX2* piRNA production. Our data revealed that in the absence of *Kdm3*, dozens of gene-containing regions become *bona fide* germinal dual strand piRNA clusters. Indeed, they produce piRNAs originating from both DNA strands, become transcribed in a Moonshiner-dependent manner and enriched in di-and tri-methylation of lysine 9 of histone H3 (H3K9me2/3) and in Rhino, an HP1-like protein. Eggs laid by *Kdm3* GLKD females do not hatch and show developmental defects phenocopying loss of function of genes included into the new piRNA clusters, suggesting an inheritance of functional ovarian “auto-immune” piRNAs. Our results demonstrate that some gene-containing regions are actively prevented for piRNA production by proteins that counteract piRNA cluster emergence. Hence, a non-piRNA-producing state is therefore not a “by default” state but rather a cellular lock that is actively controlled for some genomic loci.

**KEY FACTS:**

1. Hfp regulates the expression of Kdm3 *via* its splicing
2. Kdm3 prevents genomic regions containing coding genes from becoming piRNA clusters
3. Embryos from Kdm3 mutant females show developmental phenotypes suggesting that auto-immune piRNAs are functional and alter the expression of genes embedded in newly established piRNA clusters

---

How piRNA clusters are defined and activated at each generation remains elusive: repetitive DNA sequences, specific chromatin marks, flanking transcription units and maternal inheritance of homologous piRNAs are all features that appear critical for their determination ^6–9^. Using *BX2*, a locus made of seven *P-lacZ-white* (*P(lacW)*) transgenes and behaving as a dual-strand piRNA cluster ^7^, we previously showed that an environmental stress can activate *de novo BX2* likely through an increase of its antisense transcription ^10^. Using the bimodal epigenetic state ability of *BX2*, that can be in an ON or an OFF state for piRNA production, we designed a genetic screen allowing us to test the effect of germ line knockdown (GLKD) of genes on *BX2* activation (Fig. 1a,b). We tested 491 small hairpin RNA (shRNA) producing lines ^11^ affecting 341 candidate genes mainly involved in RNA biology and chromatin remodeling (Extended Data Table 1). Among those, only two independent lines enabled the conversion of *BX2^OFF^* into *BX2^ON^*, as shown by their ability to functionally repress a homologous euchromatic *P(lacZ)* target (Fig. 1c-g). The first line is designed to specifically target the splicing factor *half pint (hfp)* and the second line targets specifically *Kdm3. Kdm3* encodes a JmjC domain-containing histone demethylase, the single *Drosophila* orthologue of mammalian KDM3A and KDM3B that catalyze H3K9me2 demethylation ^12^. KDM3B is involved in gene activation in leukemia cells ^13–15^. In Drosophila, Kdm3 was found in substantially increased association with H3K9 lysine-to-methionine-containing mono-nucleosomes ^16^. In addition, its loss-of-function has been reported to act as an enhancer of variegation ^17^ while its overexpression acts as a suppressor of variegation ^16^ suggesting that Kdm3 acts as a structural antagonist to heterochromatin.

**Figure 1.**
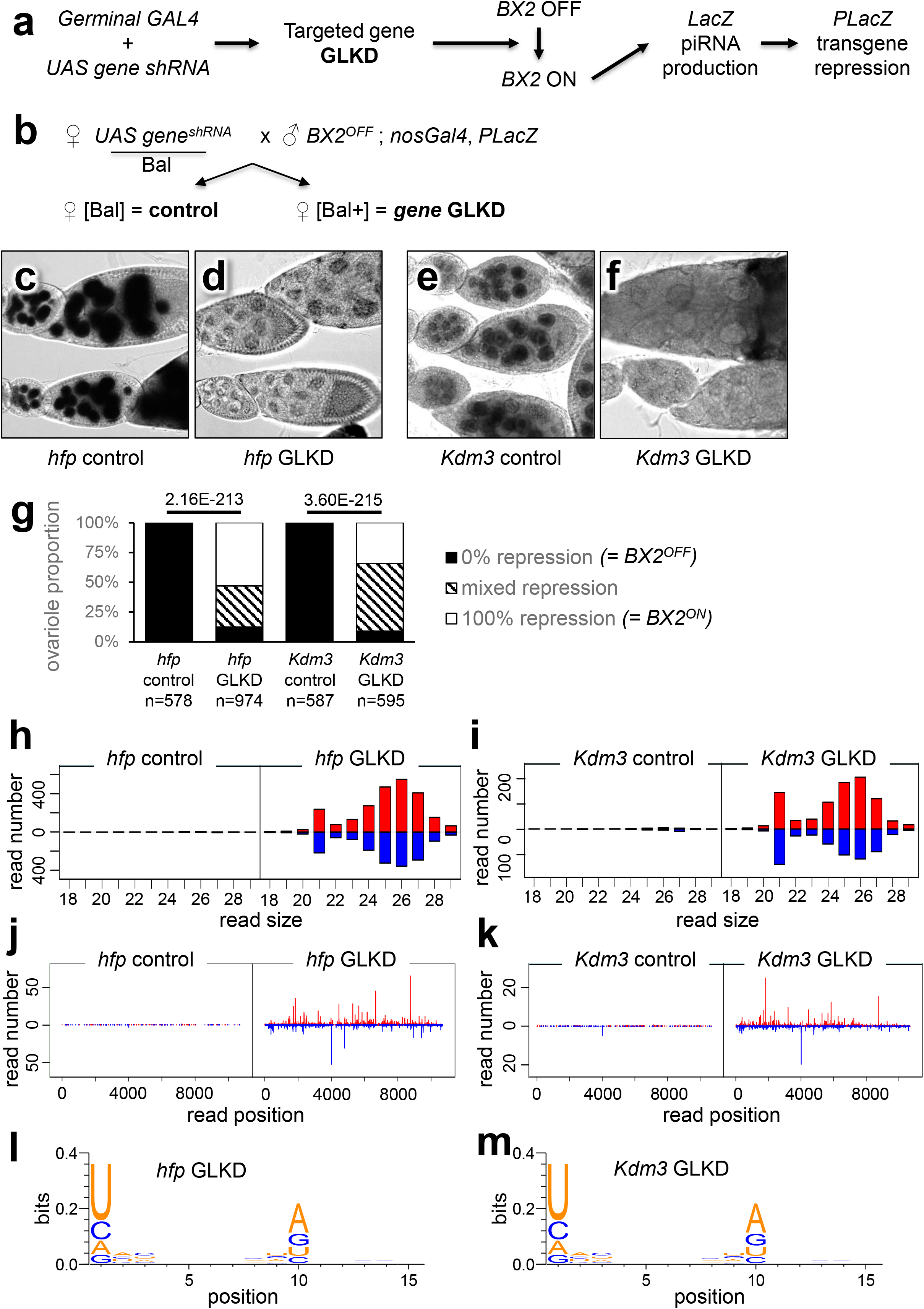
*hfp* and *Kdm3* GLKD lead to *de novo* activation of the *BX2* piRNA cluster. **a**, Strategy followed to identify genes required to maintain *BX2* in an inactive state (*BX2^OFF^*). **b**, Transgenic females carrying an *UAS gene^shRNA^* (small hairpin RNA that targets a specific gene) were crossed with males carrying the *BX2^OFF^* cluster, a transgene expressing GAL4 under the *nanos* promoter that is germline specific (*nosGAL4*) and an euchromatic *P(lacZ)* transgene that expresses the *lacZ* gene in the female germline. *P(lacZ)* is used as a readout for the epigenetic state of the *BX2* locus. When *BX2* is OFF (*BX2^OFF^*) no *LacZ* piRNAs are synthesized and the *LacZ* gene from the euchromatic *P(lacZ)* is expressed (ovarioles with stained nuclei in egg chambers after X-gal staining). When *BX2* is ON (*BX2^ON^*), *LacZ* piRNAs are synthesized, inducing *P(lacZ)* transgene silencing (ovarioles with unstained nuclei in egg chambers after X-gal staining). Control sisters allows determination if *BX2* remains stably OFF in the absence of GLKD. **c**, *BX2* remains OFF in the absence of *hfp* GLKD. **d**, *hfp* GLKD in the presence of *BX2* led to the repression of *P(lacZ)*. **e**, *BX2* remains OFF in the absence of *Kdm3* GLKD. **f**, *Kdm3* GLKD in the presence of *BX2* led to the repression of *P(lacZ)*. **g**, Ovariole repression was measured by counting the number of ovarioles showing no, partial or complete repression of the *P(lacZ)* target among egg chambers. p-values are indicated above bars (Pearson’s Chi-squared test). n = number of ovarioles counted. **h-i**, Comparison of the size distribution of 18 to 29 nt ovarian small RNAs matching *BX2* sequences between *BX2* control (*BX2^OFF^*) and *BX2* activated by *hfp* GLKD (**h**) or by *Kdm3* GLKD (**i**). Positive and negative values correspond to sense (red) and antisense (blue) reads, respectively. **j-k**, Comparison of 23 to 29 nt unique mappers along the *BX2* sequence between *BX2* control (*BX2^OFF^*) and *BX2* activated by *hfp* GLKD (**j**) or *Kdm3* GLKD (**k**). **l-m**, Logo showing the enrichment in a U at the first position, 66.8% 1U, n=28029 reads in *hfp* GLKD ovaries (**l**) and 72.0% 1U, n=11806 reads in *Kdm3* GLKD ovaries (**m**) and the enrichment of an A at the tenth position among the paired reads (49.3% 10A, n=2237 for *hfp* GLKD (**l**) and 57.7% 10A, n=929 for *Kdm3* GLKD (**m**).

GLKD effects were confirmed with characterized *hfp* and *Kdm3* mutant alleles. While amorphic or strong alleles of *hfp* lead to homozygous embryonic lethality, *hfp^9^* and *hfp^13^* are two hypomorphic alleles that allow adult survival ^18^. Different allelic mutant combinations led to the conversion of *BX2^OFF^* into *BX2^ON^*, confirming the specific role of *hfp* in this process (Extended Data Fig. 1a-f). Similarly, a *trans*-heterozygous combination of null alleles of *Kdm3* ^17,19^ led to the *BX2* conversion, confirming the role of *Kdm3* in the *BX2^OFF^* state maintenance (Extended Data Fig. 1g-i). Whole genome sequencing analyzes of small RNAs extracted from ovaries of controls, *hfp* and *Kdm3* GLKD ovaries revealed that, in agreement with the functional silencing of the *P(lacZ)* target, a bulk of 23 to 29 nucleotides small RNAs was produced all along the *BX2* sequence in both *hfp* and *Kdm3* GLKD ovaries when compared to their respective control sisters (Fig. 1h-k, Extended Data Table 2). These small RNAs were enriched in a U at 5’ end and present, among the paired reads, an enrichment of an A at the tenth position, known as the ping-pong signature (Fig. 1i-j, ^3^). Taken together, our results show that *hfp* or *Kdm3* GLKD induces piRNA production and *trans*-silencing capacities of the *BX2* locus, demonstrating for the first time that a locus made of repeated sequences could require a genetically active process to maintain a non-piRNA producing state (OFF).

## Stability of *BX2* conversion through generations

Interestingly, *BX2 hfp* GLKD flies are fertile, allowing us to check the stability of the *BX2* OFF to ON conversion in following generations and after a return to a wild type dosage of *hfp* (Extended Data Fig. 2a). Ovarian small RNA analyses revealed that the progeny of *BX2 hfp* GLKD flies that do not produce shRNA against *hfp* anymore, however maintained the production of numerous piRNAs matching *BX2* sequence when compared to the progeny of *BX2* control flies (Extended Data Fig. 2b-d). These piRNAs are enriched in U at the 5’ end and enrichment of an A at the tenth position among the paired reads (Extended Data Fig. 2e). The *BX2^ON^* state was fully maintained in seven independent lines established from seven single G2 *BX2* females and tested during 20 generations (Extended Data Table 3). These results demonstrate that, once established by a transient reduction of the *hfp* dosage, the acquired *BX2^ON^* state is maintained through generations likely by maternal inheritance of *BX2* piRNAs ^7,20^. In contrast to *hfp* GLKD flies, *Kdm3* GLKD and *Kdm3* KO flies do not produce viable progeny preventing us to analyze the stability of this conversion in the subsequent generations.

## Mutants of *otu* do not activate *BX2*

To further investigate the mechanism of *BX2* conversion by *hfp* GLKD, we first checked whether the direct down regulation of a known target of Hfp could activate *BX2^OFF^*. The *ovarian tumor* gene (*otu*) encodes a deubiquitinase involved in several processes including germ cell development. *Otu* sounded like a good candidate because the specific 104 kDa isoform, which is produced through Hfp-mediated splicing ^18^, includes a TUDOR domain that is shared by several proteins involved in piRNA biology (Spn-E, Krimp, Qin, Tej, Tapas, Vret, Tudor, SoYb, BoYb, Yb) and conserved in most animals ^2^. Therefore, we tested the effect of *otu* GLKD or *otu* mutant alleles on *BX2^OFF^* conversion. The *otu* GLKD context leads to atrophic gonads with few egg chambers and no *BX2* activation (n=15 females). Consistently, two allelic mutant combinations of *otu* also did not induce *BX2* conversion (Extended Data Fig. 1j). Taken together, these results demonstrate that the *hfp* GLKD effect on *BX2* conversion does not rely on *otu* splicing regulation nor on the 104 kDa isoform production.

## Hfp regulates *Kdm3* splicing

Based on the annotated Drosophila genome, *Kdm3* shares its 5’ UTR with the *CG8176* gene that encodes a F-BAR domain-containing protein potentially involved in cytokinesis (Fig. 2a, ^21^). This peculiar genomic position suggests that *Kdm3* expression relies on specific splicing factors. To test if the production of *Kdm3* transcripts depends on *hfp*, we measured by RT-qPCR the steady-state level of the two *Kdm3* isoform RNAs in different *hfp* mutant or GLKD backgrounds. Four different primer sets revealed that both *Kdm3* RA and RB spliced forms are strongly affected in all tested *hfp* mutant and GLKD backgrounds while unspliced isoforms were largely unaffected (Fig. 2b-e). To investigate whether Hfp plays a direct role on *Kdm3* splicing, RNA immunoprecipitation (RIP) experiments were carried out using Hfp-GFP expressing flies and anti-GFP antibodies followed by RT-qPCR analyses. As expected, RNA from known targets of *hfp* such as *tra2* ^22^ and *otu* ^18^ were significantly enriched when compared to a control gene *(eEF5)* (Fig. 2f). Importantly, *Kdm3* RNA was also significantly enriched, supporting the idea that *hfp* is directly involved in *Kdm3* splicing (Fig. 2f). Interestingly, we did not detect any interaction between Hfp and *BX2* transcripts, arguing against a direct effect of Hfp on *BX2* conversion (Fig. 2f).

**Figure 2.**
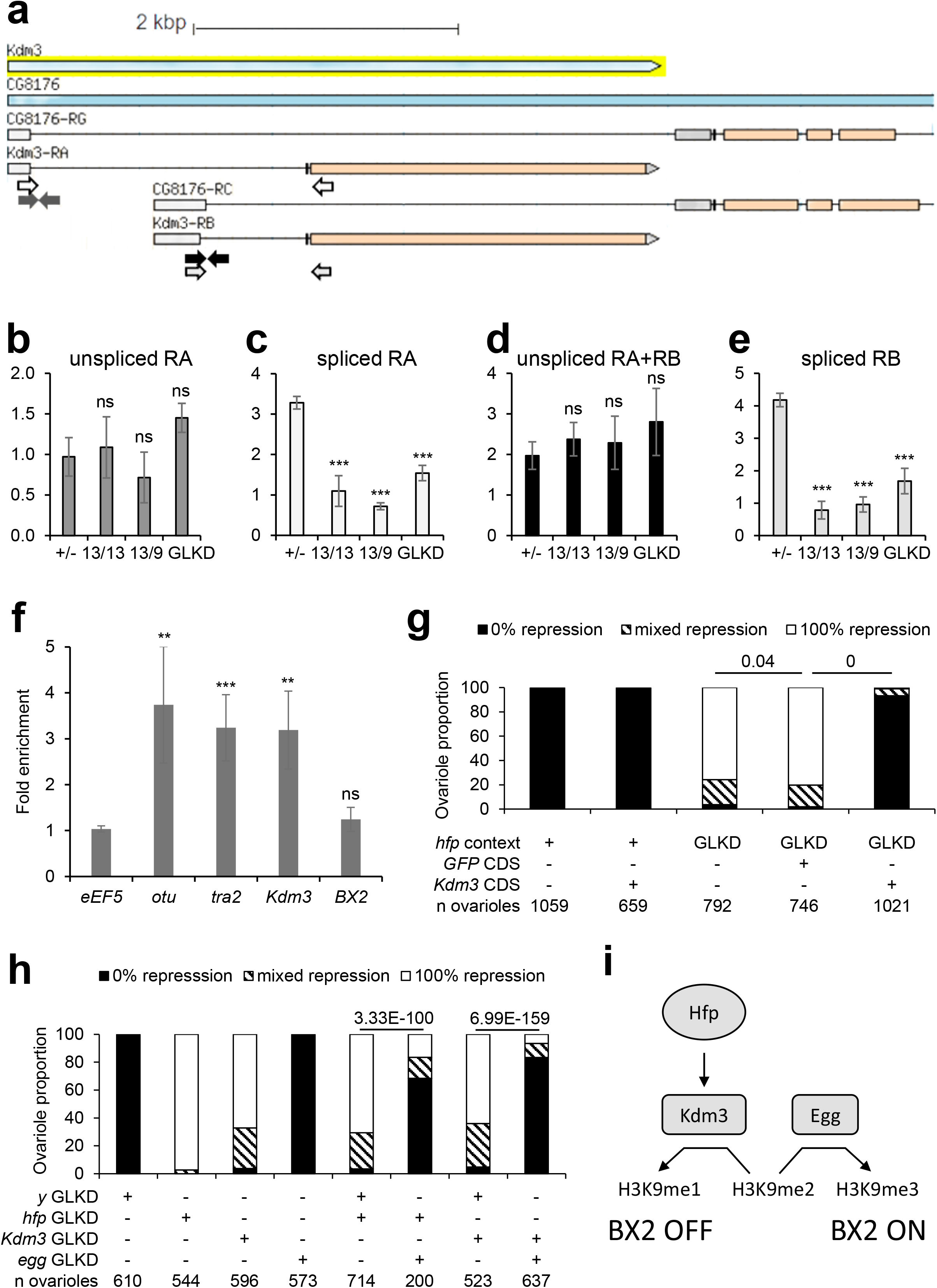
Interactions between *hfp, Kdm3* and *egg* in *BX2* conversion. **a**, The two *Kdm3* RNA isoforms (*RA* and *RB*) share a 5’UTR (light gray boxes) with two RNA isoforms of *CG8176*, suggesting a splicing control for Kdm3 production. Arrows indicate primers used for RT-qPCR experiments shown in 3b-e (colors used for arrows and histograms correspond). **b-e**, RT-qPCR experiments using primers shown in 3a revealed that both *RA* and *RB* spliced *Kdm3* isoforms are strongly affected in *hfp* mutant or GLKD contexts while unspliced isoforms are not. p-values are indicated as ns for p-value > 0.05, * for p-value < 0.05, ** for p-value < 0.01 and *** for p-value < 0001 (two sided t-test). **f**, RNA immunoprecipitation (RIP) experiment using a *hfp-GFP* construct showed that *Kdm3* transcripts behave similar to known direct targets of Hfp (*otu* and *tra2*) rather than a control gene *(eEF5). BX2* transcripts do not appear to be targeted by Hfp. **g**, Rescue experiment using Kdm3 CDS. Histogram shows the percentage of ovarioles showing homogeneous β-Galactosidase staining in all egg chambers (0% repression, *BX2^OFF^*), homogeneous non-staining in all egg chambers (100% repression, *BX2^ON^*) or mixed repression (presence of stained and unstained nuclei among egg chambers in the same ovariole). For all crosses, males bear *BX2^OFF^, nosgal4* and *P(lacZ)* transgenes and females carry a combination of *UAS-hfp shRNA* (*hfp* GLKD), *UASpRFP* (= mock CDS, as control of a potential GAL4 dilution effect on two UAS transgenes) and/or *UASpKdm3* CDS transgenes. p-values are indicated above bars (Pearson’s Chi-squared test). n = number of counted ovarioles. **h**, Epistatic relationships between *hfp, Kdm3* and *egg* GLKD upon the activation of a paternally inherited *BX2*. Histogram shows the percentage of ovarioles as in **g**. For all crosses, males bear *BX2^OFF^, nosgal4* and *P(lacZ)* transgenes while females carry a combination of *UAS-hfp shRNA, UAS-Kdm3 shRNA, UAS-egg shRNA* and/or *UAS-y shRNA* as a mock for a potential GAL4 dilution effect on two UAS transgenes. n = number of counted ovarioles., **i**, Model of regulation of the epigenetic state of *BX2* by modulation of the methylated state of the lysine 9 of histone H3 (H3K9) by Kdm3 and Egg.

## *Kdm3* and *egg* act antagonistically on the *BX2* epigenetic state

To test whether the *BX2* conversion in the *hfp* GLKD background functionally depends on *Kdm3*, we constructed a transgene containing the coding sequence (CDS) of *Kdm3* whose expression relies on the GAL4/UAS system (see material and methods). Using a driver line expressing GAL4 in the germline, we showed that the germinal expression of *Kdm3* CDS prevents the *BX2* conversion in an *hfp* GLKD background (Fig. 2g). Indeed, in that context, 93% of ovarioles (n=1021) showed β-Galactosidase expression in all egg chambers, thus demonstrating that *BX2* was not converted into an active piRNA cluster. This result strongly suggests that *Kdm3* splicing performed by Hfp is critical for *BX2* conversion. We then hypothesized that *BX2* is actively maintained OFF by Kdm3, a H3K9me2 demethylase enzyme that might counteract the function of an antagonist protein involved in the piRNA pathway to avoid piRNA cluster conversion. One of the best candidates is Eggless/SetDB1 (Egg), an H3K9 methyltransferase shown to be required for germinal piRNA cluster determination ^5,23,24^. We tested this hypothesis by analyses of the epistasis relationship between *egg*, *hfp* and *Kdm3* GLKD. The concomitant loss of *egg* and *hfp* on the one hand or of *egg* and *Kdm3* on the other hand prevents *BX2* conversion (Fig. 2h). These results show that *egg* is required for the *BX2* epigenetic conversion in a *hfp* or *Kdm3* GLKD background, very likely through control of the H3K9 methylation level (Fig. 2i).

## Other transgenic clusters can be activated upon *hfp* or *Kdm3* GLKD

Previously, we have shown that the stability throughout generations of *P(lacW)* clusters activated by maternally inherited homologous piRNAs (paramutation) depends on the size of the cluster, *i.e*. the number of transgenes making the cluster ^7^. Here, we tested whether the number of *P(lacW)* transgenes present in the same locus influences the conversion efficiency in *hfp* or *Kdm3* GLKD backgrounds (Extended Data Fig. 3a-b). As observed for the paramutation phenomenon, the longer the cluster, the higher the conversion rate (Extended Data Fig. 3c). Furthermore, smaller clusters (1 and 2 copies) could not be activated for piRNA production revealing that the conversion requires a minimum of four repeats and does not depend on the genomic integration location. Next, we asked whether a subtelomeric piRNA cluster could be stimulated for piRNA production in *hfp* or *Kdm3* GLKD backgrounds. Previously, we have described that the activation for piRNA production of transgenes embedded in subtelomeric piRNA clusters was spontaneous but delayed when the transgenes were paternally inherited ^25^. Indeed, four generations were required to reach a full production of piRNAs and a full repression of a *P(lacZ)* target in a β-Galactosidase functional assay ^25^. Here, we show that *hfp* and *Kdm3* GLKD dramatically increased subtelomeric transgenes activation during the first generation (Extended Data Fig. 3d,e). These results indicate that the activation effect is not restricted to the *BX2* piRNA cluster family. As the subtelomeric transgenes are embedded into natural piRNA clusters, these finding also suggest that Kdm3 may counteract the propagation of piRNA from neighboring piRNA clusters, thereby helping in defining the frontiers of piRNA clusters.

## Dozens of genomic regions become piRNA clusters upon *Kdm3* GLKD

To know whether other genomic regions could be actively turned ON for piRNA production, we performed a comparative analysis of the sequencing of three replicates of control and *Kdm3* GLKD ovarian small RNA libraries. Despite its strong effect on the conversion of *BX2* family and subtelomeric piRNA clusters, *Kdm3* GLKD did not induce major changes in the global piRNA production in ovaries. The number of 23-29 nt RNAs that map onto the Drosophila genome was not significantly modified while a slight decrease could be observed for 23-29 nt RNAs matching previously defined piRNA clusters (Table 1, ^3,26^). On the opposite, a slight increase could be observed regarding transposon sequences, particularly when unique mappers were considered (Table 1). Taken together, these observations revealed that the *Kdm3* GLKD context did not modify the global amount of piRNAs but could partly change their genomic sources.

**Table 1.** Number of 23-29 nt small RNA matching Drosophila genome, previously defined piRNA clusters and transposon sequences. Total or unique mappers are indicated. n is the mean number of reads given in read per million (rpm). sd is the standard deviation observed between the three biological replicates. p-value from t-test is given.

|  | Control |  | *Kdm3* GLKD |  | p-value (t-test) |
| --- | --- | --- | --- | --- | --- |
|  | n (rpm) | sd | n (rpm) | sd |  |
| Drosophila genome (total) | 256539 | 3987 | 269805 | 8524 | 0.07108124 |
| Drosophila genome (unique) | 60577 | 891 | 64494 | 2783 | 0.08098947 |
| piRNA clusters defined in (Brennecke et al. 2007) (unique) | 29894 | 799 | 27405 | 428 | 0.00891689 |
| piRNA clusters defined in (Chen et al. 2021) (unique) | 33969 | 1078 | 32193 | 839 | 0.02126752 |
| Transposons (total) | 174551 | 5708 | 189898 | 6001 | 0.03260395 |
| Transposons (unique) | 10477 | 251 | 11874 | 340 | 0.0046077 |

To question the source of piRNA production, the Drosophila genome was first split into 1 kb bins and the unique 23-29 nt reads isolated from control and *Kdm3* GLKD ovaries were then mapped onto these bins (Extended Data Table 2). Bins producing a differential amount of 23-29 nt RNAs (DEseq2 analysis, fold change ≥8, *p* value <0.001) were recovered. Selected bins separated from less than 3 kb were grouped (clusterization) resulting in 134 differentially 23-29 nt-expressing regions (Extended Data Table 4). They range from 6 to 182 kb and represent a total of 2.5 Mbp. Eleven intergenic regions (totaling 144 kbp) produce less 23-29nt RNAs in *Kdm3* GLKD than in control while 123 regions (totaling 2.38 Mbp) produces significantly more 23-29 nt RNA in *Kdm3* GLKD than in control. We identified 13 regions with a high mean amount of 23-29 nt RNAs (MA plot red dots in Fig. 3a) including *white* (*w*) sequences (R100) corresponding to *BX2* (*P(lacW)* transgenes) and *jing* (R4), the gene flanking the *42AB* locus, the largest natural germline piRNA cluster in *D. melanogaster*. Other selected regions contain genes coding mostly for transcriptional factors implicated in embryonic development (Fig. 3a). Small RNAs produced by these regions originate from both strands (Fig. 3b), the 23-29 nt RNAs present an enrichment in U at 5’ end (63.5% 1U, n=98560) and A at the tenth position among the paired reads (56% 10A, n=1397, Fig. 3c). Taken together, these data suggest that these 23-29 nt RNAs are *bona fide* piRNAs. RNA-seq experiments performed on total ovarian RNA followed by a DEseq2 analysis between *Kdm3* GLKD and control revealed an increase in the steady-state RNA level of the genes embedded in those regions producing *de novo* piRNAs (Extended Data Table 2, Fig. 3d). Strikingly, in wild type conditions, most of the genes contained in these regions are not transcribed in ovaries. RT-qPCR experiments confirmed increased RNA levels in the *Kdm3* GLKD context and further showed that the piRNAs depend on the transcription factor Moonshiner (Moon) (Fig. 3e). Moon has been characterized as part of a rhino-dependent transcription machinery that enables the initiation of transcription of germline piRNA clusters ^9^. Taken together, these results strongly support the idea that these regions have become genuine double strand piRNA clusters.

**Figure 3.**
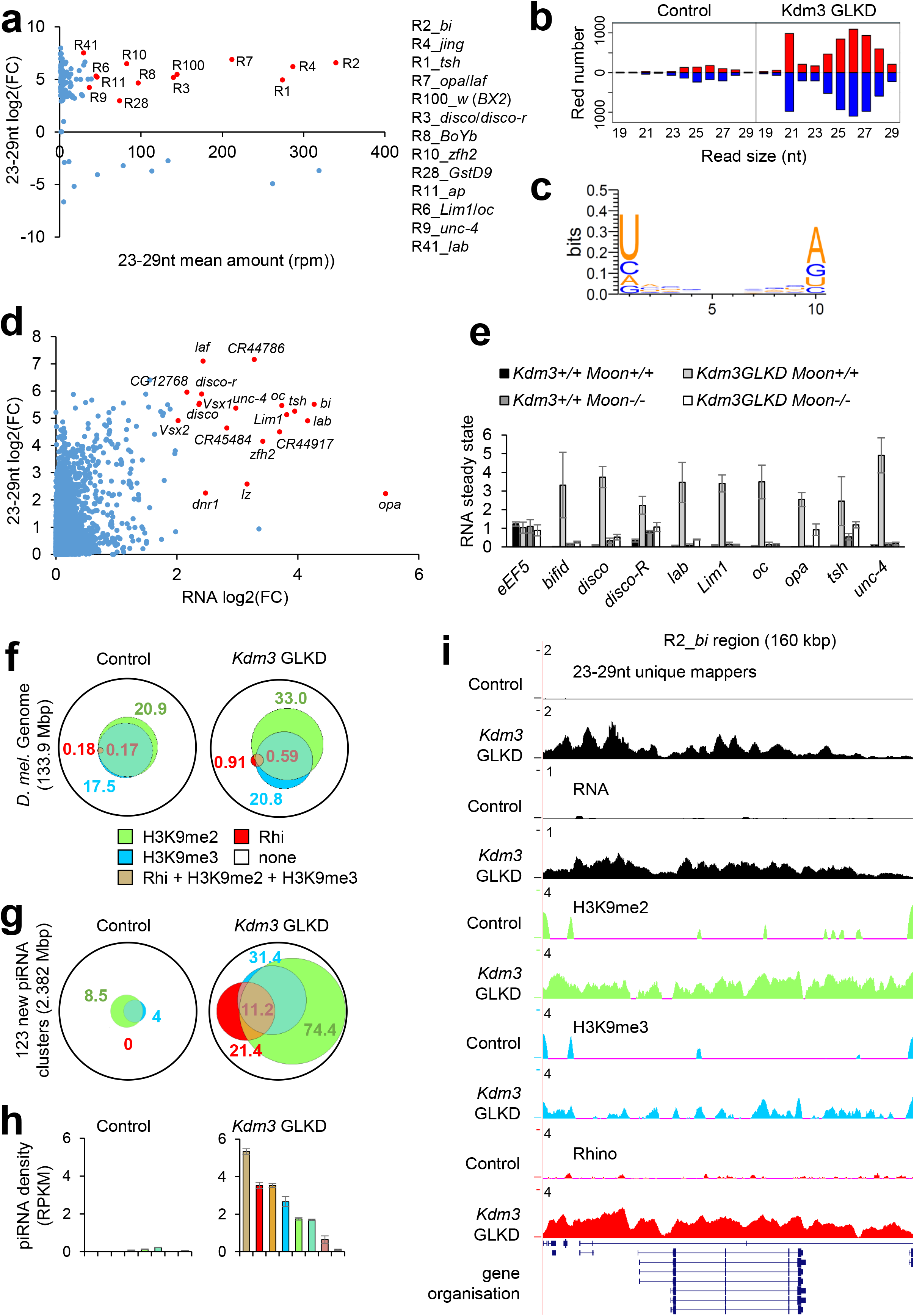
Characterization of new piRNA clusters upon *Kdm3* GLKD. **a**, Ovarian **s**mall RNAs (23-29 nt, unique mappers per 1 kb bins) fold change (log2, y axis) between *Kdm3* GLKD and control compared to the mean amount of 23-29 nt (x-axis) revealed that 123 gene-containing regions produce a significant increased amount of small RNAs. Thirteen major regions are depicted in red and listed with included gene names. **b**, Comparison of the size distribution of small RNAs (18-29 nt) matching the 13 major producing regions (red numbers in Fig. 3a) between control and *Kdm3* GLKD contexts. Positive and negative values correspond to sense (red) and antisense (blue) reads, respectively. **c**, Logo showing the enrichment in U1 and A10 of the 10 nt paired small RNAs that match the 13 major producing regions in *Kdm3* GLKD. **d**, Ovarian small RNAs (23-29 nt, unique mappers on genes) fold change (log2, y axis) between *Kdm3* GLKD and control compared to RNA fold change between *Kdm3* GLKD and control (log2, x-axis) from total RNA-seq shows that genes embedded in the new piRNA producing loci also produce more transcripts. **e**, RT-qPCR measurements confirm the increase of RNAs observed by RNA-seq (in **b**) in *Kdm3* GLKD (light grey columns) compared to control (dark columns). We show here that this increase depends on Moonshiner (white columns). The loss of Moonshiner itself does not alter the RNA steady state level of these regions (dark grey columns). **f**, Venn diagram of ChIP-seq analyses revealing genome proportions that are enriched in H2K9me2, H3K9me3 and/or Rhino in control or *Kdm3* GLKD. **g**, Venn diagram showing the proportion of H3K9me2, H3K9me3 and/or Rhino enrichments upon the 123 new piRNA clusters (as defined in **a**) in control and in *Kdm3* GLKD. **h**, Density of 23-29 nt unique mappers (in RPKM) by the 123 regions according to their chromatin state in control and in *Kdm3* GLKD. **i**, Summary maps of small RNA-seq (23-29 nt) unique mappers, RNA seq, ChIP seqs on R2_*bi* region. Y-axis values are in rpm.

## Chromatin prefigures piRNA production

We therefore determined H3K9me2, H3K9me3 and Rhino (Rhi) occupancy through chromatin immunoprecipitation followed by sequencing (ChIP-seq) from control and *Kdm3* GLKD ovaries (Extended Data Table 2). MACS2 analyses revealed that in control ovaries, 20.9% of the genome is enriched in H3K9me2 and 17.5% in H3K9me3, which almost completely overlap (97.5% of H3K9me3-enriched regions are also H3K9me2-enriched, Fig. 3f, Extended Data Table 5). Rhi-enriched regions represent only 0.18% of the genome (~244 kbp), a proportion compatible with previous observations (^27^, Fig. 3f, Extended Data Table 5). 0.17% of the genome is co-enriched in Rhi, H3K9me2 and H3K9me3, meaning that 93.3% of the Rhi-enriched regions are co-enriched with H3K9me2 and H3K9me3 (Fig. 3f, Extended Data Table 5). This strong overlap suggests a functional relationship between the H3K9 methylation level and Rhi. In *Kdm3* GLKD ovaries, 33% of the genome is enriched in H3K9me2, likely as the direct consequence of the Kdm3 H3K9 demethylase depletion (Fig. 3f). This increase in H3K9me2 also affects the H3K9me3 distribution, since a larger part of the genome is now enriched in H3K9me3 (20.8%) but the proportion overlapping with H3K9me2-enriched regions remains stable (Fig. 3f, Extended Data Table 5). More dramatically, the genome proportion on which Rhi binds is five times greater in *Kdm3* GLKD (0.91% vs 0.18%). 0.59% of the genome (~790 kbp) became co-enriched in Rhi, H3K9me2 and H3K9me3, representing 63.5% of Rhino-enriched regions (Fig. 3f and Extended Data Table 5). Considering the 2,382 kbp of sequences corresponding to the 123 new piRNA clusters determined through differential production of 23-29 nt in *Kdm3* GLKD *vs* control (Fig.3a), we noticed that, in the control context, they were poorly enriched in H3K9me2 (8.5%) or H3K9me3 (4%) and that none of them were enriched in Rhi (Fig. 3g and Extended Data Table 6). In *Kdm3* GLKD background, the chromatin landscape of these regions is drastically modified since 74.4% of the sequences were enriched in H3K9me2, 31.4% in H3K9me3 and 21.4% in Rhi. The proportion of sequences enriched in all of the three marks reached 11.2% (~268 kbp). Finally, we calculated the piRNA density for each subtype of enrichment and observed that the density is higher for regions that are co-enriched with Rhi, H3K9me2 and H3K9me3 (5.34 piRNA per kbp and per million of reads (RPKM), Fig. 3h and Extended Data Table 7). This result highlights the crucial relationship between these chromatin modifications and the piRNA production. Genomic analyses were summarized and illustrated for the *R2_bi* region (Fig. 3i) and other regions (Extended Data Fig. 4). Taken together, our results show that Kdm3 counteracts piRNA cluster determination and piRNA production from several specific gene-containing regions, thus revealing that a molecular control based on chromatin state rather than a by-default state exists to maintain some coding sequences in a piRNA non-producing state.

## Inheritance of auto-immune piRNAs may cause developmental defects in the offspring

Finally, we addressed whether these new ovarian piRNAs might affect offspring. The hatching rate of the embryos laid by *Kdm3* GLKD females was severely impaired, whatever their genotype, thus revealing a strong maternal effect (Fig. 4a). A double GLKD of *Kdm3* and *egg* in females dramatically increased the hatching rate of their progeny (Fig. 4a). This result reinforces the hypothesis that the developmental issues of the embryos laid by the *Kdm3* GLKD females are linked to the emergence of the new piRNA producing regions. Embryos issued from *Kdm3* GLKD females stopped their development at different stages, 17% (n=369) reaching the first larval stage (L1) without hatching. The analysis of their cuticular phenotypes revealed defects in the mouth formation (Fig. 4c) as well as in the denticle belts (Fig. 4f) when compared to controls (Fig. 4b, e). These phenotypes are reminiscent of *labial* (*lab*) and *occeliless (oc*, previously known as *orthodenticle, otd)* loss of function, respectively (Fig. 4d,g, ^28,29^). Interestingly, *lab* and *oc* are included in the new piRNA producing regions defined in *Kdm3* GLKD ovaries (Fig. 3). RT-qPCR experiments performed on 3-5 h embryos revealed that *lab* and *oc*, as well as most of the genes embedded in these new piRNA clusters, were less expressed in *Kdm3* GLKD female progeny when compared to control progeny (Fig. 4h). Taken together, these data suggest that new piRNAs produced in the ovaries of *Kdm3* GLKD females are inherited in the progeny and impaired the expression of several corresponding embryonic developmental genes. As piRNAs are often compared to a genomic immune system, we propose to name these new piRNAs, “auto-immune piRNAs”.

**Figure 4.**
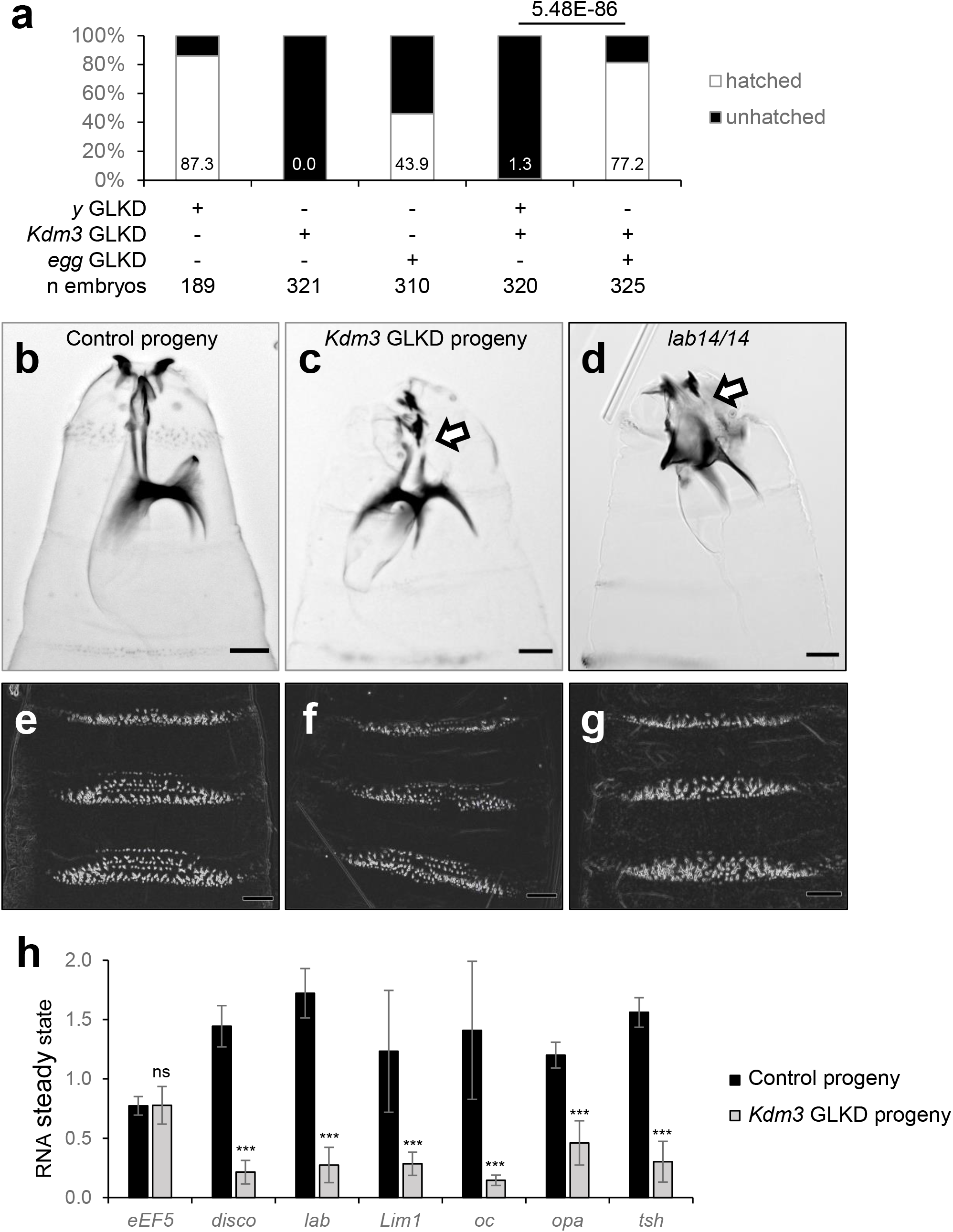
The progeny of *Kdm3* GLKD females display strong maternal phenotypes. **a**, As indicated above each column in percentage, the hatching rate of the *Kdm3* GLKD progeny is strongly impaired when compared to control (*mock* GLKD). Double *Kdm3* and *egg* GLKD almost fully rescues the hatching rate. **b-c**, Sclerotized structures of the head of unhatched embryos issued from control females (**b**) or *Kdm3* GLKD females (**c**). The lack of continuity (as shown by an arrow) between the sclerites in the anterior part and the more posterior structures (ventral plate and arms) is even stronger in a *labial* complete loss-of-function embryo (homozygous for *lab*^14^) (**d**). Scale bar is 30 μm. **e-g**, Ventral cuticular pattern of the embryos (a1 to a3 denticle bands) reveals the loss of denticle rows in the anterior region of the abdomen in the *Kdm3* GLKD progeny (**f**) compared to control (**e**). This phenotype is reminiscent of *oc/otd* loss of function (**g**). **h**, RT-qPCR measurements of transcripts in embryos arising from *Kdm3* GLKD females reveal that upregulated genes embedded into deregulated regions in the mothers’ ovaries (Fig 3) are downregulated in their progeny.

To conclude, we determined that HFP controls splicing of the histone demethylase *Kdm3* gene and that *Kdm3* down regulation increases the H3K9me2 chromatin enrichment by 50%. A subset of these regions then become enriched in H3K9me3 probably partly through the action of the histone methyltransferase Eggless. Then, a more restricted genomic region subset is enriched in Rhino, leading to the formation of new piRNA clusters and to the production of auto-immune piRNAs in ovaries, which impair the development of the progeny likely through their maternal inheritance (Fig. 5). Indeed, our results suggest that chromatin composition prefigures the piRNA cluster determination in ovaries. Therefore, the balance between the protection of the genome against transposable elements and appropriate developmental gene expression is primarily defined in the female germline and then transmitted to the offspring. This balance is dependent on tight control that defines genomic regions through chromatin modifications that regulate piRNA production.

**Figure 5.**
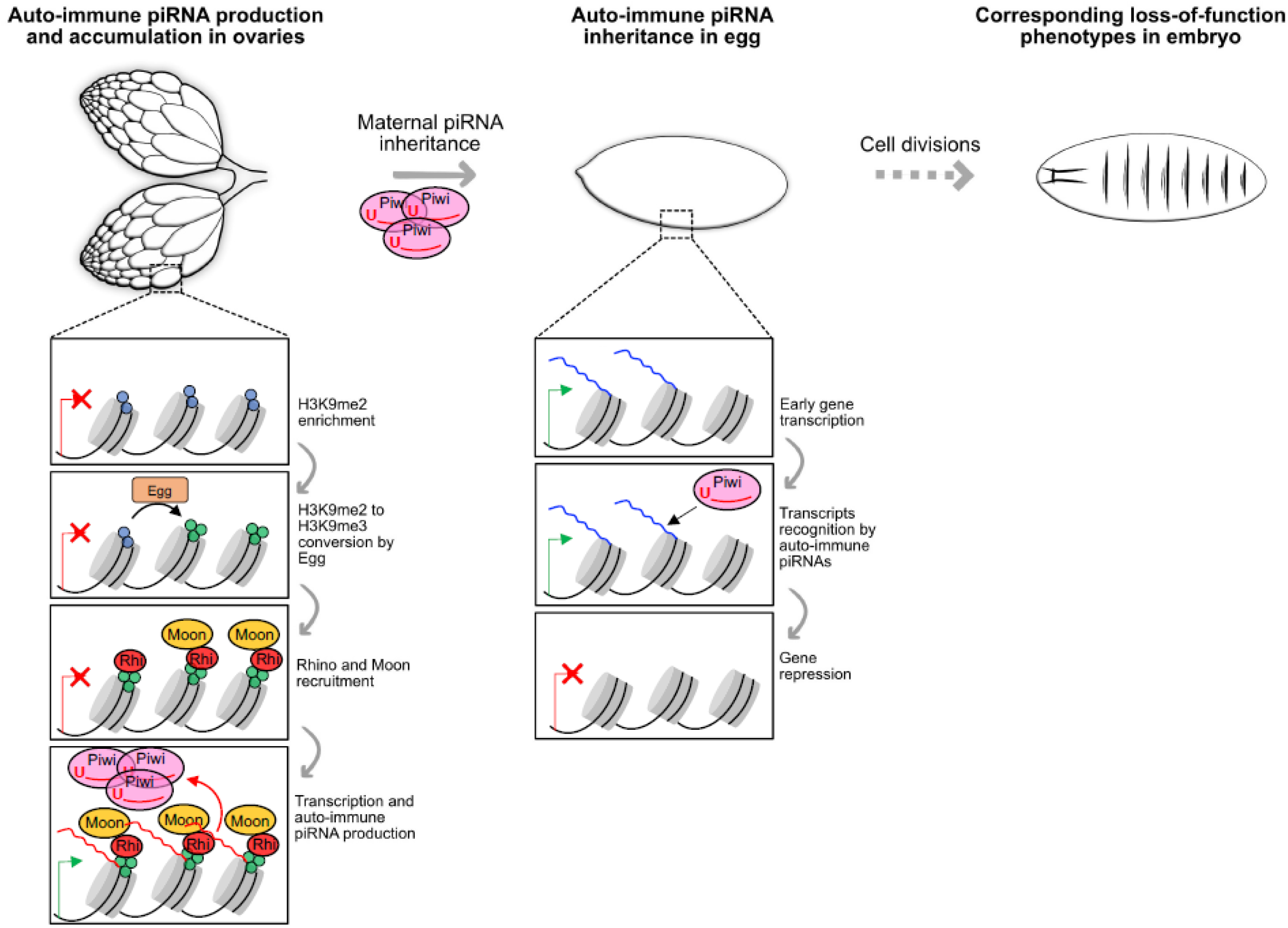
Model proposing that the maternal inheritance of the piRNAs produced by gene-coding regions in the *Kdm3* GLKD ovaries leads to embryonic developmental phenotypes due to a loss of function of the same genes in the progeny.

## EXTENDED DATA

**Extended Data Figure 1. *hfp* and *Kdm3* mutation leads to the *de novo* activation of the *BX2* piRNA cluster. a-e,** X-gal staining of ovaries. Genotypes are given. *hfp^9^* and *hfp^13^* are two hypomorphic alleles allowing adult survival ^18^. **f**, Ovariole repression was measured by counting the number of ovarioles showing no, partial or complete repression of the *P(lacZ)* target among egg chambers. Interestingly, we observed that the strongest hypomorphic allele (*hfp^9^*) leads to the strongest *BX2* activation (almost 100% in homozygous state) while the weakest allele (*hfp^13^*) leads to a weakest *BX2* activation. Heterozygous combination leads to an intermediate *BX2* activation. n = number of counted ovarioles. **g-h**, X-gal staining of ovaries. Genotypes indicated correspond to allelic combination of *Kdm3* loss-of-function alleles: a heterozygous *Kdm3^KO/+^* does not lead to the activation of *BX2* (**g**) while a *trans*-heterozygous loss-of-function combination leads to *BX2* conversion (**h**). **i**, Ovariole repression was measured as previously. n = number of counted ovarioles. **j**, Combinations of *otu* mutant alleles do not induce *BX2* conversion. The parental cross corresponds to *otu^11^* / *M5; P(lacZ)* females crossed by *otu^pat^; BX2^OFF^ / CyO* males. *otu^pat^* can be *otu^11^*, an EMS induced C343Y substitution in the TUDOR domain and hence specifically affecting the activity of the 104 kDa Otu isoform ^30^, or *otu^7^*, an EMS induced nonsense mutation K424@ (or K382@ depending on the isoform), leading to truncated non-functional proteins. Whatever the allelic combination, *BX2* never switched on.

**Extended Data Figure 2. Stability of the *hfp* GLKD mediated *BX2* conversion at the next generation. a**, Genetic crosses performed to analyze the progeny of control or *hfp* GLKD females. **b**, Comparison of the size distribution of ovarian small RNAs (18-29 nt) matching *BX2* sequences between progeny of control and *hfp* GLKD. Positive and negative values correspond to sense (red) and antisense (blue) reads, respectively. **c**, Comparison of unique 23-29 nt mappers along the *BX2* sequence between progeny of control and *hfp* GLKD. **d**, Comparison of multiple 18-29 mappers along the *hfp* sequence between *hfp* GLKD flies (G1) and their progeny (G2) shows that G2 flies no longer produce shRNA against *hfp* 3’ UTR (as indicated by the black arrow). **e**, Logo showing the enrichment in U1 and A10 of the 10 nt paired small RNAs that match *BX2* in the *hfp* GLKD progeny. These piRNAs are enriched in U at position 1 (67.7% n=34609 reads) and exhibit a ping-pong signature as an enrichment of 10A among paired reads (47.4% n=2421).

**Extended Data Figure 3. Other allelic piRNA clusters are activated upon *Kdm3* GLKD. a**, Origin and number of *P(lacW)* copies in each studied cluster ^7,31,32^. All clusters are inserted at the same locus. **b**, Each cluster was paternally inherited in order to measure its conversion rate in the ovaries of the progeny. The conversion was measured as the ability of the cluster to silence the *P(lacZ)* transgene. **c**, Histogram showing the conversion rate of the six clusters in three genetic backgrounds, *white (w)* GLKD serving as control, *hfp* GLKD or *Kdm3* GLKD. n = number of counted ovarioles. **d**, Following a similar scheme, telomeric transgenes (*P-1152*) were paternally inherited and their conversion rate was analyzed in the ovaries in the progeny as in **c**. **e**, Histogram showing the conversion rate of telomeric transgenes that are paternally inherited in the three different genetic contexts. n = the ovariole number.

**Extended Data Figure 4.** Panels summarizing *Kdm3* GLKD induced modifications of 23-29 nt production, RNAseq and ChIPseq analyses on several genomic regions including the *jing*, *lab*, *disco-r/disco* and *tsh* genes.

**Extended Data Table 1. List of tested shRNA lines.** Gene name, gene symbol, CG number, Bloomington stock number, gonad phenotype and progeny viability are indicated.

**Extended Data Table 2. Summary of small RNAseq, RNAseq and ChIPseq data.** Name of library, genotype and depth are given. The fastq.gz files were deposited on the GEO under the number GSE203279.

**Extended Data Table 3. Stability of *hfp* GLKD induced *BX2* conversion through subsequent generations.** A complete and stable conversion was observed for seven lines established from seven single G2 *BX2* females and tested during 20 generations by cross with *P(lacZ)* males and ß-Galactosidase assay of the progeny ovaries. n = number of tested females.

**Extended Data Table 4. Determination of regions presenting a differential expression of unique 23-29 nt RNA.** Coordinates, size, rank, gene content, ID, mean 23-29 nt expression, sigma, basemean and fold change are given. nc: not calculable.

**Extended Data Table 5. Genome wide ChIP-seq analyses using H3K9me2, H3K9me3 and Rhino antibodies.** MACS2 analyses gave the number of significant enriched peaks for both H3K9 marks and Rhino compared to their respective input. The total length of enriched sequences and their proportion in relation to the total size of the *Drosophila melanogaster* genome (%) are given. BED intersect analyses allowed to determine overlap coordinates and to define co-enriched regions.

**Extended Data Table 6. Chromatin enrichment of the 123 new piRNA clusters in control and *Kdm3* GLKD contexts.** MACS2 analyses revealed the presence of H3K9me2, H3K9me3 and/or Rhino enrichments upon the 123 new piRNA producing regions in *Kdm3* GLKD and in these regions in the control condition. The cumulative size of these regions is given.

**Extended Data Table 7. piRNA production linked to the chromatin state of the 123 new piRNA clusters.** The 23-29 nt (unique mappers) production was obtained using ovarian small RNAseq analyses and the rpm value was calculated using normalization factors estimated from each library depth.

## Materials and Methods

### Transgenes and strains

All transgenes are in the *w^1118^* background. The *P(lacW)* transgene (FBtp0000204) is composed of 5’ and 3’ *P* element extremities (*P5* and *P3*, respectively), *E. coli lacZ* gene *(lacZ), D. melanogaster miniwhite* gene (*W*) and backbone plasmid pBR322 sequence (pBR). The *BX2* line (FBti0016766) carries seven *P(lacW)*, inserted in tandem and in the same orientation at cytological site 50C on the second chromosome ^31^. The transgene insertion site is located in an intron of the *AGO1* gene ^7^. Homozygous individuals are rare and sterile and the stock is maintained in heterozygous state with a *Cy*-marked balancer chromosome. ß-Galactosidase activity from these transgenes is not detected in the germline. *P(lacZ)* corresponds to *BQ16* (FBti0003435) expressing *lacZ* only in the germline from *P(A92)* (FBtp0000154), a *P-lacZ-rosy* enhancer-trap transgene located on the third chromosome. Homozygous flies are viable. Another *P(lacZ)* located on the second chromosome at 60B7 was used when necessary, a *P(PZ)* transgene (FBtp0000210) corresponding to a *P-lacZ-rosy* enhancer-trap transgene and expressing ß-Galactosidase in the germline and somatic cells of the female gonads (Bloomington stock number *11039* (FBst0011039). Homozygous flies are not viable and the stock is maintained over a *Cy*-marked balancer chromosome. The *P-1152* line carries two *P(lArB), P-lacZ-rosy* enhancer-trap transgenes (FBti0005700) that are inserted into subtelomeric sequences of the *X*chromosome ^33^. The *nosGAL4* transgene mainly used is from *w[1118]; P{w[+mC]=GAL4::VP16-nos.UTR}CG6325[MVD1]* line (FBti0012410). The *nosGAL4* from the *w[*]; PBac{w[+mW.hs]=GreenEye.nosGAL4}Dmel6* line (FBti0131635) was also occasionally used and gave the same results. *otu* alleles are *otu^11^*, an EMS induced C343Y substitution in the TUDOR domain and hence specifically affecting the activity of the 104 kDa Otu isoform ^30^ or *otu^7^*, an EMS induced nonsense mutation K424@ (or K382@ depending on the isoform), each leading to truncated non-functional proteins. *Kdm3* alleles are *Kdm3^KO^*, a gift of Dr Michael Buszczak, ^17^ and *Kdm3^MI13382^*, a MiMic transgene inserted in the JmJc domain that inactivates the gene, Bloomington stock center #59126, ^19^. Small hairpin RNAs (shRNAs) are listed in Extended Data Table 1. *hfp* and *Kdm3* shRNA that allow *BX2* activation are TRiP line #34785 and TRiP line #32975, respectively. *egg* shRNA showing epistasis interactions with *hfp* and *Kdm3* shRNA is TRiP line #36797. Mock shRNA are *y* shRNA TRiP line #64527 on chromosome 2 or *w* shRNA TRiP line #33644 on chromosome 3. Additional information about stocks are available upon request or at Flybase: “http://flybase.bio.indiana.edu/”. Flies were grown on a corn-based medium with agar. All crosses were performed at 25°C.

### Cloning procedures

*EGFP* sequences were amplified by PCR from *pGEM5Z(+)-EGFP* (Addgene #65206) and were cloned into *p(UASp)* (FBtp0010350) at the *BamHI* site. *Kdm3* coding sequence was amplified from wild type genomic DNA (*CantonS*) and cloned into *p(UASp-EGFP)* at the *Eco*RI site in frame with the *EGFP* sequence. Sequence integrity was verified by sequencing. Map and sequence are available on request. Plasmid *p(UASp-Kdm3-EGFP)* was then injected into *w^1118^* embryos by BestGene Inc. Several independent lines were recovered. An insertion on the *X* chromosome was used for rescue experiments.

### β-Galactosidase staining

Ovarian *lacZ* expression assays were carried out using X-gal (5-bromo-4-chloro-3-indolyl-beta-D-galactopyranoside) overnight staining at 37°C as previously described ^34^, except that ovaries were fixed afterwards for 10 min. After mounting in glycerol/ethanol (50/50), ovarioles were counted as fully repressed if all egg chambers were white (100% repression), non-repressed if all egg chambers were blue (0% repression) or mixed if some egg chambers were blue while others were white (mixed repression). Images were acquired with an Axio-ApoTome (Zeiss) and ZEN2 software.

### Cuticle preparation

Experiment has been done as previously described ^35^. Laid embryos were allowed to age for 24 h at 25°C. They were rinsed in water and then dechorionated using 8% sodium hypochlorite solution for 2 min. Embryos were rinsed again in water and the vitelline membrane was removed mechanically using a needle. Water was replaced by 1:1 lactic acid:Hoyer’s-based medium. Embryos were transferred onto a glass microscope slide and placed under a cover slip. Slides were incubated at 60°C overnight. Phase contrast images were obtained using the x20 objective of an Axio-ApoTome (Zeiss) and ZEN2 software.

### RNA extraction

For ovaries samples, 10 to 20 pairs of ovaries were manually dissected in 1X PBS. For embryo samples, 15 to 20 μl of embryos aged of 3-5 h after egg laying were collected, washed with water and flash frozen in liquid nitrogen. Total RNA was extracted using TRIzol (Life Technologies) as described in the reagent manual (http://tools.lifetechnologies.com/content/sfs/manuals/trizol_reagent.pdf). For the RNA precipitation step, 100% ethanol was used instead of isopropanol and washes were made in 80% ethanol instead of 75% ethanol. Up to six biological replicates were used for each genotype.

### RT-qPCR experiments

For each sample, 2 μg of total RNA was treated with DNase (Fermentas). For RT-qPCR experiments, 1 μg of DNase-treated RNA was used for reverse transcription using random hexamer primers (Fermentas). Real-time qPCR was performed on triplicates of each sample. *RpL32* was used as reference. The same series of dilution of a mix of different RT preparations was used to normalize the quantity of transcripts in all RT preparations leading to standard quantity (Sq) values. Variations between technical triplicates was very low when compared to variations between biological replicates. The mean of the three technical replicates was then systematically used (meanSq). For each biological sample, we calculated the ratio meanSq(gene)/meanSq(*RpL32*) to normalize the transcript quantity. Then, the mean of each sample ratio was used to compare the two conditions.

### Native RNA-IP followed by RT-qPCR

For each sample, 80 pairs of ovaries from well-fed females (2 or 3 days) were manually dissected in cold 1X PBS, flash frozen in liquid nitrogen and conserved at −80°C. Ovaries from *hfp-GFP* and *GFP* negative control lines were lysed and grinded with pestle in RNA-IP lysis buffer (KCl 150 mM, Tris pH7.4 25 mM, EDTA 5 mM, DTT 0.5 mM, Nonidet P-40 0.5%), freshly supplemented with Protease Inhibitor Cocktail (1 tablet per 10 ml of RNA-IP lysis buffer) and RNase Inhibitor (40 U per 1 ml of RNA-IP buffer). Lysates were sonicated using Bioruptor (Bioruptor Standard Diagenode) for 7 minutes (15 sec ON, 60 sec OFF) and cleared by centrifugation at 12,000 rpm for 15 min at 4°C. 10% of cleared lysate was set aside to serve as input samples and the remainder was incubated at 4°C with anti-GFP antibodies (Roche #11814460001) for 3 h under gentle rotation. Magnetic beads coupled to G protein (Dynabeads # 10003D, Invitrogen) were washed two times with RNA-IP lysis buffer, transferred into immunoprecipitated lysate and incubated at 4°C for 1 h under gentle rotation. The beads were washed 5 times with RNA-IP lysis buffer for 10 min under gentle rotation at 4°C. Total RNA was extracted from input and beads using TRI reagent (Sigma-Aldrich Cat #T9424) as described in reagent manual (https://www.sigmaaldrich.com/technical-documents/protocols/biology/tri-reagent.html). For RNA precipitation step, 100% ethanol was used instead of isopropanol and for wash steps 80% ethanol was used instead of 75% ethanol. For each genotype, three to four biological replicates were used. For each input sample, 2 μg of total RNA was treated with DNase (New England Biolabs) and for the RT-qPCR experiment, 1 μg of DNase-treated RNA was used for reverse transcription using random hexamer primers (Fermentas). For each immunoprecipitated sample, total RNA was treated with DNase, and total DNase-treated RNA was used for reverse transcription using random hexamer primers. Real-time PCR was performed on duplicates for each biological sample leading to Cycle threshold (Ct) values. Variations between technical duplicates were very low compared to variations between biological replicates. The mean of the two technical replicates was then systematically used (meanCt). Data have been analyzed as described in ^36^. For each sample, the IP fraction is normalized beside input to take account of sample preparation difference as follows: ΔCt [normalized RIP] = (meanCt [RIP] – (meanCt [Input] – Log2 (Input Dilution Factor))) where meanCt [RIP] is the Ct value measure for immunoprecipitated samples, meanCt [Input] is the Ct value measure for input and Input Dilution Factor corresponds to the RNA fraction set aside for input (in this experiment, 10% of RNA fraction was set aside, thus Input Dilution Factor is 10). Antibody signal specificity was confirmed by comparing *hfp-GFP* to *GFP* negative control, Fold Enrichment was calculated for each sample as follows: Fold Enrichment = 2^(-ΔCt[normalized RIP]–ΔCt [normalized NS]^.

### List of primers used in this study

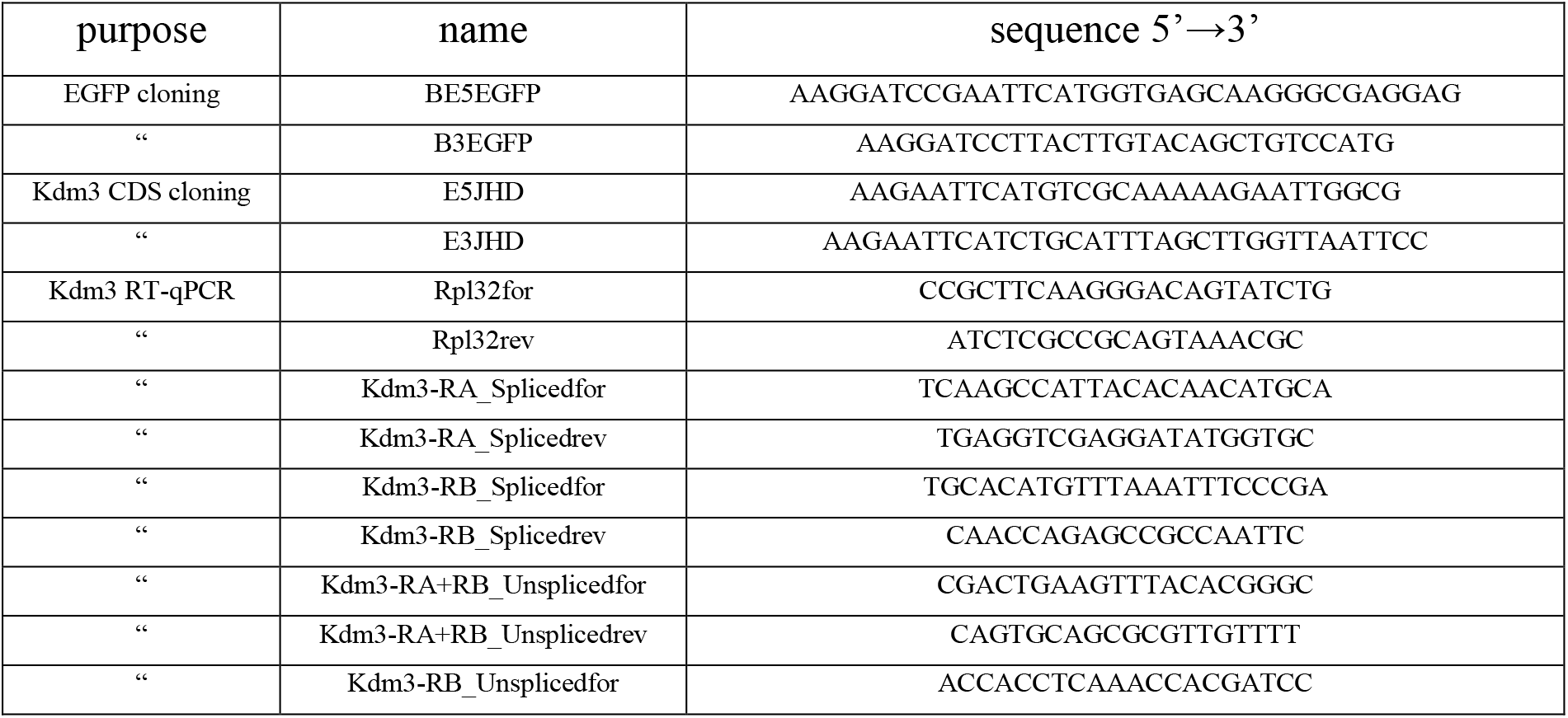
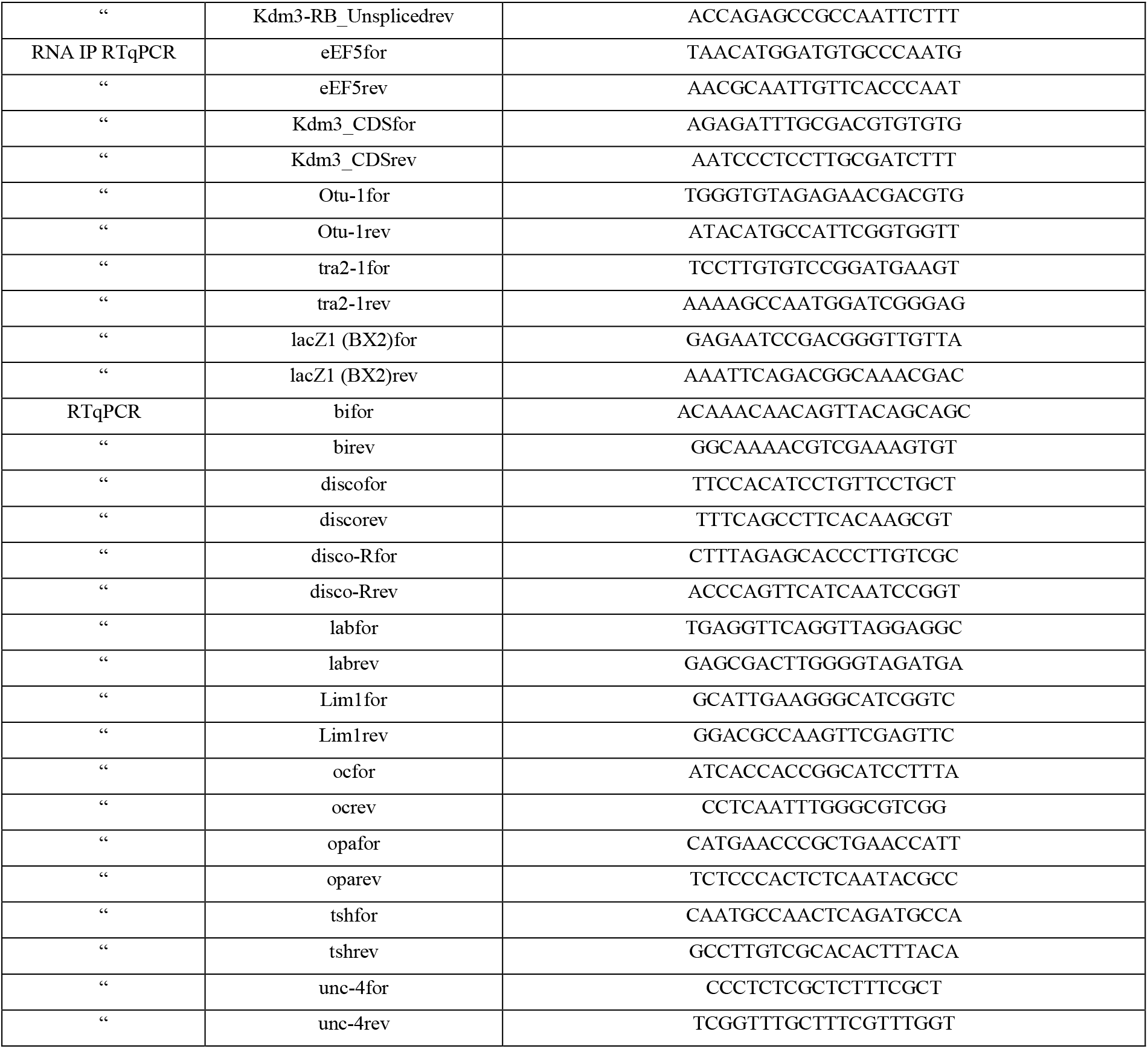

### small RNA sequencing analyses

A small RNA fraction of 15 nt to 30 nt in length was obtained following separation of total RNA extracted from dissected ovaries on a denaturing polyacrylamide gel. This fraction was used to generate multiplexed libraries with Illumina TruSeq Small RNA library preparation kits (RS-200-0012, RS200-0024, RS-200-036 or RS-200-048) at *Fasteris* (http://www.fasteris.com). A house protocol based on TruSeq, which reduces 2S RNA (30 nt) contamination in the final library, was performed. Libraries were sequenced using Illumina HiSeq 2000 and 2500. Sequence reads in fastq format were trimmed from the adapter sequence 5’-TGGAATTCTCGGGTGCCAAG-3’ and matched to the *D. melanogaster* genome release 6 (dm6) using Bowtie (Table 1, Extended Data Table 2, ^37^). For subsequent analyses, we used a cleaned version of the genome in which Y, U and mitochondrial chromosomes were removed. This version, named “dm6_clean”, has been deposited at the GEO under accession number GSE203279.

Sequence length distributions, small RNA mapping and small RNA overlap signatures were generated from bowtie alignments using Python and R (http://www.r-project.org/) scripts, which were wrapped and run in Galaxy instance publicly from ARTbio platform available at http://mississippi.fr. Tools and workflows used in this study may be downloaded from this Galaxy instance. For library comparisons, read counts were normalized to one million reads (Extended Data Table 2). For small RNA mapping (Figure 1, 3, Extended Data Figure 2), we took into account only 23-29 nt RNA reads that uniquely aligned to reference sequences (unique mappers). Logos were calculated using Weblogo ^38^ from 3’ trimmed reads (23 nt long) matching *P(lacW)* (Figure 1l-m) or selected genomic regions (Figure 3c). The percentage of reads containing a “U” at the first position was calculated with all 23-29 nt RNA matching the reference sequence. Distributions of piRNA overlaps were computed as first described in ^39^ and detailed in ^40^. Thus, for each sequencing dataset, we collected all of the 23-29 nt RNA reads matching *P(lacW)* whose 5’ ends overlapped with another 23-29 nt RNA read on the opposite strand. Then, for each possible overlap of 1 to 29 nt, the number of read pairs was counted. The percentage of reads containing an “A” at the tenth position was calculated within the paired 23-29 nt RNA matching the reference sequence as described in ^7^. Small RNA sequences and projects have been deposited at the GEO under accession number GSE203279.

### RNAseq analysis

Total RNA was extracted with TRIzol from hand dissected ovaries in 1X PBS and 1 μg was treated with DNase (Fermentas). After an initial quality control, libraries were prepared using the RNA RiboZero Stranded protocol by Fasteris. Indexed adapters were ligated and multiplexed sequencing was performed using Illumina HiSeq 2000 and 2500 (Extended Data Table 7). A DeSEq2 analyze on all *D. mel*. genes (release 6.36) was performed using the three biological replicates for each genotype, control or *Kdm3* GLKD. The original file has been deposited at the GEO under accession number GSE203279.

### ChIP-seq analyses

Chromatin immunoprecipitation (ChIP) was performed as previously described ^41^ with minor modifications. Briefly, 100 ovary pairs were manually dissected into Schneider media and cross-linked in 1% formaldehyde/PBS for 10 min at room temperature with agitation. The cross-linking reaction was quenched by STOP buffer (PBS 1X, Triton 0.1%, Glycine 1 M) and ovaries were washed in PBS and homogenized in a glass douncer: first slightly dounced in PBST 0.1% and centrifugated 1 min 400g, followed by strong douncing in cell lysis buffer buffer (KCL 85 mM, HEPES 5 mM, NP-40 0.5%, Sodium butyrate 10 mM, EDTA free protease inhibitor cocktail Sigma) following by 5 min centrifugation at 2000g. We performed 2 washes with cell lysis buffer. The homogenates were then lysed on ice for 30 min in nucleus lysis buffer (HEPES 50 mM, EDTA 10 mM, N lauryl sarkosyl 0.5%, sodium butyrate 10 mM, EDTA free protease inhibitor cocktail Sigma). DNA was sheared using a Bioruptor pico from Diagenode for 10 cycles (30 sec on, 30 sec off). The sonicated lysates were cleared by centrifugation and then incubated overnight at 4°C with anti-Rhino antibody. Then 40 μL of Protein A Dynabeads was then added and allowed to bind antibody complexes by incubation for 1 h at 4°C. For H3K9me2 and me3 ChIP, 50 μL of Protein A Dynabeads were first coated with the anti-H3K9me3 or me2 antibodies and then incubated with the chromatin overnight at 4°C. Following four washing steps with high salt buffer (Tris pH 7.5 50 mM, NaCl 500 mM, Triton 0.25%, NP-40 0.5%, BSA 0.5%, EDTA pH 7.5 5 mM), DNA-protein complexes were eluted and de-cross-linked 10 h at 65°C. RNA and protein was digested by RNase A and Proteinase K treatments, respectively, before purification using Phenol/Chloroform protocol. Barcoded libraries were prepared using Illumina technology, which were sequenced on a NextSeq High (Illumina) by Fasteris for the ChIP H3K9me3 and H3K9me2 and by Jean Perrin facility for the ChIP Rhino (Extended Data Table 7).

### Data analyses

For alignments we used a cleaned version of r6.36 version of *Drosophila melanogaster* genome (dm6, Flybase) in which sequences corresponding to unmapped (chrU), Y chromosome (chrY) and mitochondrial chromosome (chrM) were removed (dm6_clean).

For small RNA-seq analyses, we first discarded the reads matching tRNA (dmel-all-tRNA-r6.30 Flybase), miRNA (dmel-all-miRNA-r6.30 Flybase), miscRNA (dmel-all-miscRNA-r6.30 Flybase) and rRNA (dm3-rRNA-sequences Galaxy Tutorial) using sR_Bowtie -m3 (Galaxy Version 2.1.1). Then, unique mapper reads were obtained using sR_Bowtie -m0 (Galaxy Version 2.1.1) and the dm6_clean genome as reference. 23-29 nt reads were aligned on 124 new piRNA production loci with sR_Bowtie -m0 (Galaxy Version 2.1.1). The reads were counted with Bamparse (Galaxy Version 3.0.0). Then the counts were normalized in reads per million (rpm), and means and ratio were calculated to plot the log2FC (*Kdm3* GLKD compared to control) on the BaseMean.

For RNA-seq analyses, reads were aligned to genes’ reference genome (all-gene-r6.36.fasta Flybase) with Bowtie_wrappers (Galaxy Version 1.2.0) and counted with featureCounts (Galaxy Version 1.6.4). The differential expression *(Kdm3* GLKD compared to control) was determined with DEseq2. The same procedure was used to identify genes with a differential production of piRNA except that the alignment was made with sR_Bowtie -m0.

For ChIP-seq analyses, reads were mapped on dm6_clean reference genome with BWA (Galaxy Version 0.7.17.4). PCR duplicates were discarded using Picard tool (Galaxy Version 2.18.2.1). Peak calling was made on mapped reads with MACS2 (Galaxy Version 2.1.1.20160309.6) using following parameters: --gsize ‘120000000’ --keep-dup ‘1’ --qvalue ‘0.05’ --nomodel --extsize ‘200’ --shift ‘0’. Narrowpeak was used for libraries obtained with Rhino antibody and broadpeak calling was used for libraries obtained with H3K9me2 and H3K9me3 antibodies ( --broad --broadcutoff=‘0.1’). To define the whole enrichment the ratio between the IP sample and Input sample was calculated. A differential analysis was also performed by calculating the ratio between IP *Kdm3* GLKD samples and IP control samples. Overlapping enriched regions were determined using bedtools intersect intervals (bedtools Galaxy Version 2.30.0). Overlaps between chromatin-marks-enriched regions and new piRNA producing regions were determined similarly. DNA sequences (in fasta format) from these overlaps were recovered from coordinates with GetFastaBed (bedtools Galaxy Version 2.30.0). Unique mappers were aligned on these sequences using sR_Bowtie -m0 to determine the piRNA density.

UCSC views were made with bigWig files generated with BamCoverage (Deeptools Galaxy Version 3.3.2.0.0) for small and long RNA-seq. Bin size was 50 bp and reads were normalized in counts per million. For Chip-seq analyses, BamCompare (Deeptools Galaxy Version 3.3.2.0.0) was used with a bin size of 100 pb and the computation of the log2 of the IP/Input ratio.

## ACKNOWLEDGEMENTS

We thank Doug Dorer, Steve Henikoff, Julius Brennecke, Michael Buszczak and the *Bloomington Stock Center* for providing stocks. We thank Julius Brennecke for providing antibodies. We thank flybase.org for providing databases. We thank Ritha Zamy for technical assistance. We thank Lori Pile for critical reading of the manuscript. We thank Christophe Antoniewski for helpful advice and development of the ARTbio platform (http://artbio.fr/).

## FUNDING

This work was supported by PhD fellowships from the *Ministère de l’Enseignement Supérieur et de la Recherche* to KC, JA and PPM and fellowships from Ligue Nationale contre le Cancer to KC. C.C. and L.T. received financial support from the CNRS, Sorbonne Université, the ANR (ANR-21-CE12-0022-01 #BiopiC) and the Association pour la Recherche contre le Cancer (Fondation ARC, PJA 20191209395). We also thank the Réseau André Picard and la Société Française de Biologie du Développement for their support on travel fellowship grants to KC. The funders had no role in study design, data collection and analysis, decision to publish, or preparation of the manuscript.

## COMPETING INTERESTS

The authors declare that no competing interest exist.

## Notes

### Competing Interest Statement

The authors have declared no competing interest.

