## Supplementary figures and images for "The histone demethylase KDM3 prevents auto-immune piRNAs production in *Drosophila*"

### Supplemental Figure 1

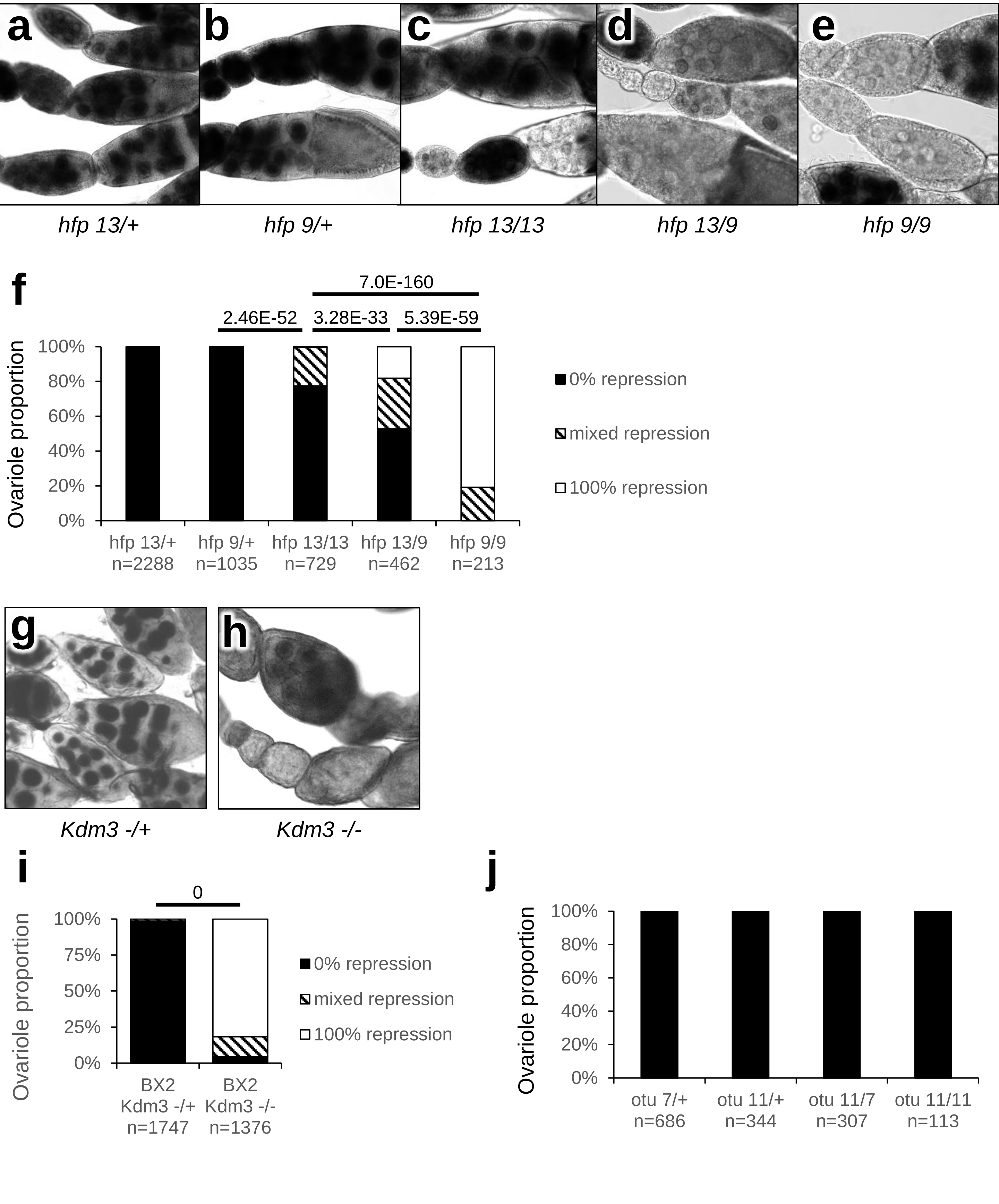


Extended Data Figure 1 Casier et al.

### Supplemental Figure 2

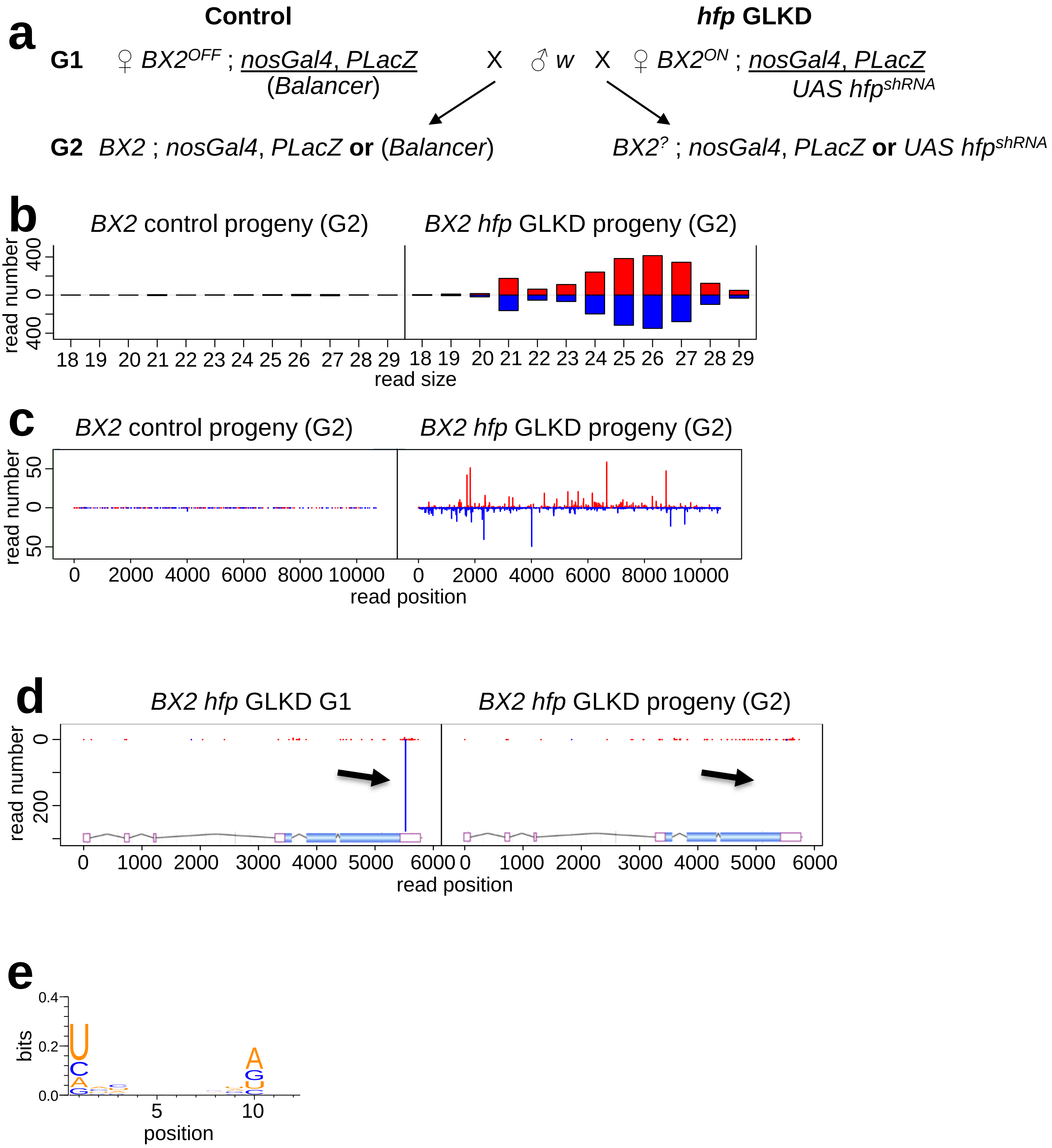


Extended Data Fig. 2 Casier et al.

### Supplemental Figure 3

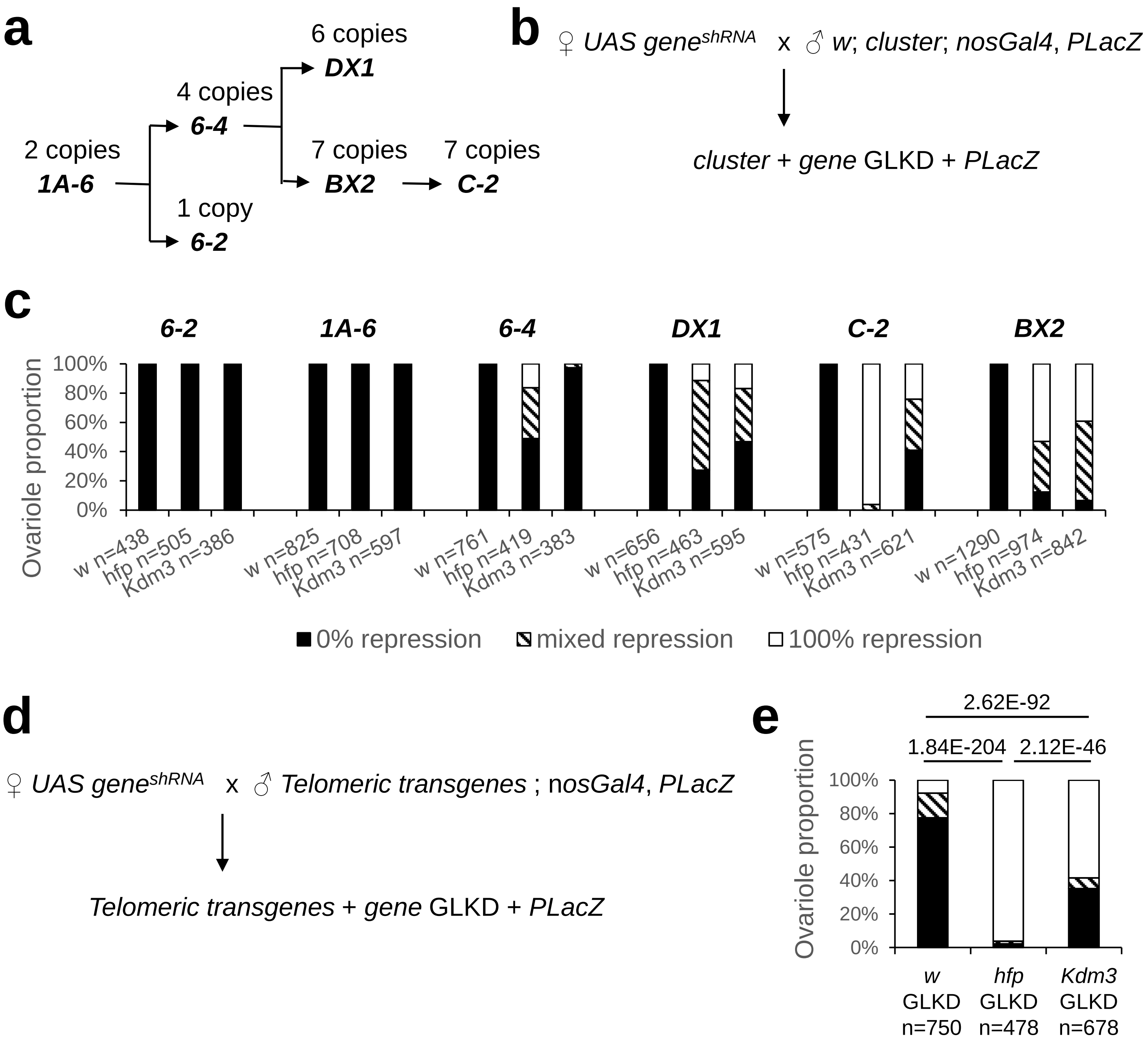


Extended data Fig. 3 Casier et al.

### Supplemental Figure 4

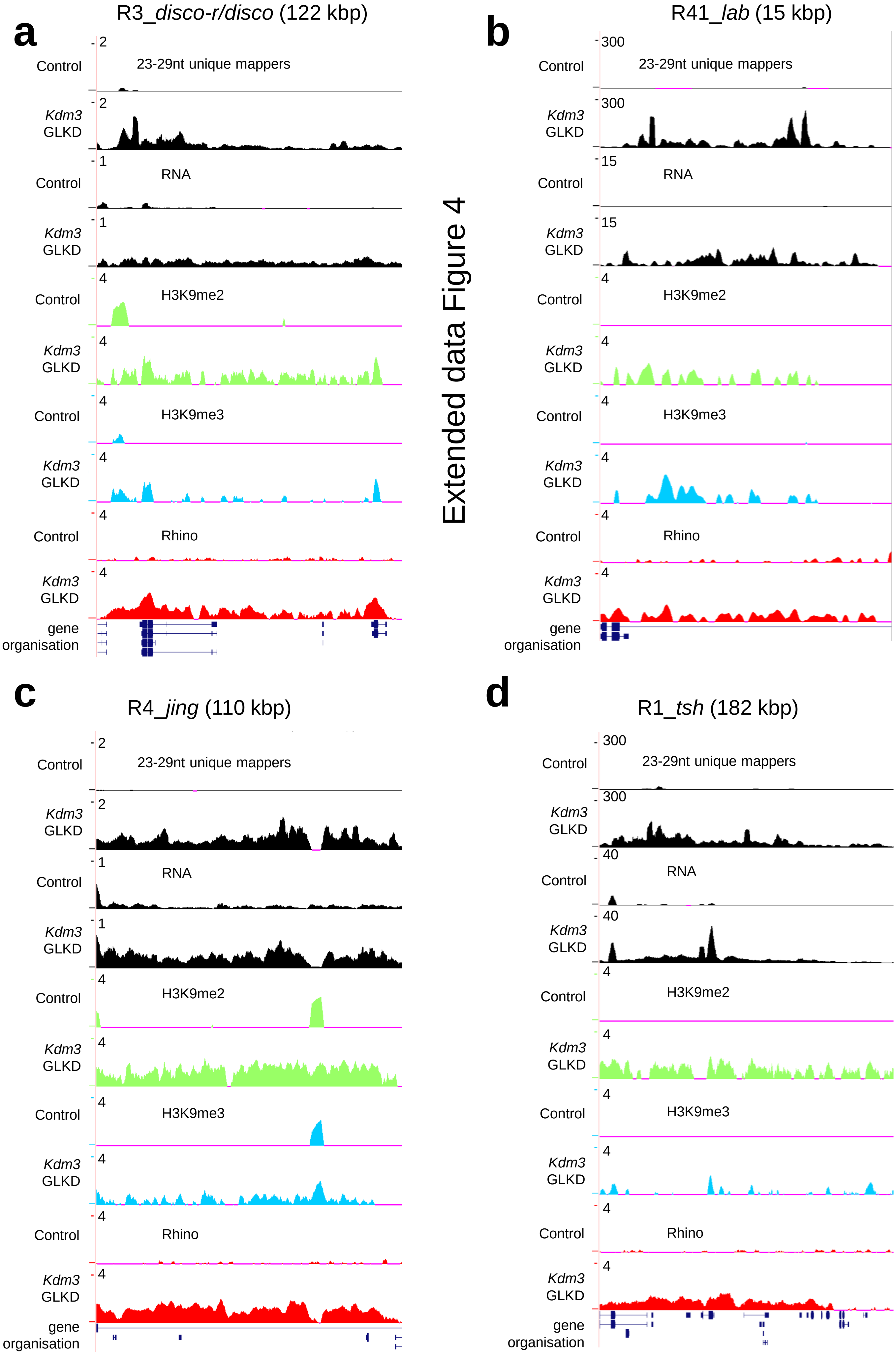


Extended data Figure 4
