## Supplemental Table 1 for "The histone demethylase KDM3 prevents auto-immune piRNAs production in *Drosophila*"

| Gene name | Gene symbol | CG | Bloomington stock | Gonad phenotype | Viable progeny |
|---|---|---|---|---|---|
| absent, small, or homeotic discs 1 | ash1 | CG8887 | 33705 | Normal | yes |
| absent, small, or homeotic discs 1 | ash1 | CG8887 | 36803 | Normal | yes |
| absent, small, or homeotic discs 2 | ash2 | CG6677 | 35388 | Normal | no |
| Ada2a-containing complex component 2 | Atac2 | CG10414 | 32890 | Normal | yes |
| Ada2a-containing complex component 2 | Atac2 | CG10414 | 53918 | Atrophy | no |
| Additional sex combs | Asx | CG8787 | 51677 | Normal | yes |
| adrift | aft | CG5032 | 63639 | Normal | yes |
| alan shepard | shep | CG32423 | 33996 | Normal | yes |
| alan shepard | shep | CG32423 | 38218 | Normal | yes |
| alan shepard | shep | CG32423 | 43545 | Normal | yes |
| Andropin | Anp | CG1361 | 55385 | Normal | yes |
| antimeros | atms | CG2503 | 12114 | Normal | yes |
| antisense RNA:CR45485 | asRNA:CR45485 | CR45485 | 62273 | Normal | yes |
| archipelago | ago | CG15010 | 34802 | Normal | yes |
| Argonaute 2 | AGO2 | CG7439 | 34799 | Normal | yes |
| Argonaute 3 | AGO3 | CG40300 | 35232 | Normal | no |
| Argonaute 3 | AGO3 | CG40300 | 44543 | Normal | no |
| Argonaute-1 | AGO1 | CG6671 | 33727 | Normal | no |
| ATP-dependent chromatin assembly factor large subunit | Acf1 | CG1966 | 35575 | Normal | yes |
| ATP-dependent chromatin assembly factor large subunit | Acf1 | CG1966 | 35575 | Normal | yes |
| Autophagy-related 1 | Atg1 | CG10967 | 44034 | Normal | yes |
| Autophagy-related 17 | Atg17 | CG1347 | 36918 | Normal | yes |
| Autophagy-related 18a | Atg18a | CG7986 | 34714 | Normal | yes |
| Autophagy-related 2 | Atg2 | CG1241 | 35177 | Normal | yes |
| Autophagy-related 8a | Atg8a | CG32672 | 34340 | Normal | yes |
| B52 | B52 | CG10851 | 37519 | Normal | yes |
| basket | bsk | CG5680 | 32977 | Normal | yes |
| basket | bsk | CG5680 | 35594 | Normal | yes |
| basket | bsk | CG5680 | 36643 | Normal | yes |
| basket | bsk | CG5680 | 53310 | Normal | yes |
| belle | bel | CG9748 | 35185 | Atrophy | no |
| belle | bel | CG9748 | 35302 | Normal | yes |
| blanks | blanks | CG10630 | 33667 | Normal | nd |
| brahma | brm | CG5942 | 34520 | Normal | yes |
| brahma | brm | CG5942 | 35210 | Normal | yes |
| brahma | brm | CG5942 | 35211 | Normal | yes |
| Brahma associated protein 60kD | Bap60 | CG4303 | 32503 | Normal | yes |
| Brahma associated protein 60kD | Bap60 | CG4303 | 33954 | Normal | yes |
| brivido-3 | brv3 | CG13762 | 33763 | Normal | yes |
| brivido-3 | brv3 | CG13762 | 36774 | Normal | yes |
| bruno1 | bru1 | CG31762 | 35394 | Atrophy | no |
| bruno1 | bru1 | CG31762 | 38983 | Atrophy | no |
| bruno1 | bru1 | CG31762 | 44483 | Normal | no |
| Caper | Caper | CG11266 | 44431 | Normal | yes |
| cap-n-collar | cnc | CG43286 | 32863 | Normal | yes |
| cap-n-collar | cnc | CG43286 | 40854 | Normal | yes |
| Carbonic anhydrase 2 | CAH2 | CG6906 | 41836 | Normal | yes |
| Carbonic anhydrase 2 | CAH2 | CG6906 | 65081 | Normal | yes |
| CG10418 | CG10418 | CG10418 | 44458 | Atrophy | no |
| CG10445 | CG10445 | CG10445 | 43137 | Normal | yes |
| CG11447 | CG11447 | CG11447 | 43207 | Normal | yes |
| CG12054 | CG12054 | CG12054 | 50511 | Normal | yes |
| CG12054 | CG12054 | CG12054 | 50910 | Normal | yes |
| CG1239 | CG1239 | CG1239 | 42825 | Normal | yes |
| CG1239 | CG1239 | CG1239 | 42912 | Atrophy | no |
| CG1239 | CG1239 | CG1239 | 44023 | Atrophy | no |
| CG1239 | CG1239 | CG1239 | 44506 | Atrophy | no |
| CG1239 | CG1239 | CG1239 | 55400 | Normal | yes |
| CG12493 | CG12493 | CG12493 | 42791 | Normal | yes |
| CG13397 | CG13397 | CG13397 | 51808 | Normal | yes |
| CG14131 | CG14131 | CG14131 | 63716 | Normal | yes |
| CG1673 | CG1673 | CG1673 | 38363 | Normal | yes |
| CG17544 | CG17544 | CG17544 | 64632 | Normal | yes |

| | | | | | |
|---|---|---|---|---|---|
| CG17724 | CG17724 | CG17724 | 35676 | Normal | yes |
| CG18537 | CG18537 | CG18537 | 53960 | Normal | yes |
| CG2926 | CG2926 | CG2926 | 34941 | Normal | nd |
| CG3036 | CG3036 | CG3036 | 43179 | Normal | yes |
| CG31075 | CG31075 | CG31075 | 50654 | Normal | yes |
| CG31075 | CG31075 | CG31075 | 62535 | Normal | yes |
| CG31262 | CG31262 | CG31262 | 53970 | Normal | yes |
| CG40006 | CG40006 | CG40006 | 34691 | Normal | yes |
| CG40160 | CG40160 | CG40160 | 44268 | Normal | yes |
| CG42542 | CG42542 | CG42542 | 35736 | Normal | yes |
| CG42588 | CG42588 | CG42588 | 63698 | Normal | yes |
| CG4267 | CG4267 | CG4267 | 32332 | Normal | yes |
| CG4267 | CG4267 | CG4267 | 56900 | Normal | yes |
| CG43373 | CG43373 | CG43373 | 64884 | Atrophy | no |
| CG4404 | CG4404 | CG4404 | 62415 | Normal | yes |
| CG4461 | CG4461 | CG4461 | 53298 | Normal | yes |
| CG5001 | CG5001 | CG5001 | 32392 | Normal | yes |
| CG5728 | CG5728 | CG5728 | 36592 | Normal | yes |
| CG6283 | CG6283 | CG6283 | 51498 | Normal | yes |
| CG6415 | CG6415 | CG6415 | 51867 | Normal | yes |
| CG7255 | CG7255 | CG7255 | 58352 | Normal | yes |
| CG7409 | CG7409 | CG7409 | 51433 | Normal | yes |
| CG7878 | CG7878 | CG7878 | 35229 | Normal | yes |
| CG8778 | CG8778 | CG8778 | 36793 | Normal | yes |
| CG9684 | CG9684 | CG9684 | 36880 | Normal | yes |
| CG9925 | CG9925 | CG9925 | 35811 | Normal | yes |
| Chitinase 2 | Cht2 | CG2054 | 35717 | Normal | no |
| Chitinase 2 | Cht2 | CG2054 | 60369 | Normal | yes |
| Chromatin assembly factor 1, p55 subunit | Caf1-55 | CG4236 | 34069 | Atrophy | no |
| Chromosome associated protein D3 | Cap-D3 | CG31989 | 36615 | Normal | yes |
| Cleavage and polyadenylation specific factor 6 | Cpsf6 | CG7185 | 34804 | Atrophy | no |
| CMP-sialic acid synthase | Csas | CG32220 | 54843 | Normal | yes |
| Complex I intermediate-associated protein, 30 kDa | CIA30 | CG7598 | 55660 | Normal | yes |
| corkscrew | csw | CG3954 | 33619 | Normal | yes |
| corkscrew | csw | CG3954 | 35215 | Normal | no |
| corkscrew | csw | CG3954 | 35638 | Normal | no |
| corkscrew | csw | CG3954 | 60448 | Normal | yes |
| crossover suppressor on 3 of Gowen | c(3)G | CG17604 | 62969 | Normal | yes |
| C-terminal Binding Protein | CtBP | CG7583 | 32889 | Normal | no |
| cup | cup | CG11181 | 35406 | Normal | no |
| Cyclic-AMP response element binding protein A | CrebA | CG7450 | 42562 | Normal | yes |
| Cyclin T | CycT | CG6292 | 32976 | Atrophy | no |
| Cyclin T | CycT | CG6292 | 35168 | Normal | yes |
| Cyclin-dependent kinase 4 | Cdk4 | CG5072 | 36060 | Normal | yes |
| Cyclin-dependent kinase 4 | Cdk4 | CG5072 | 57031 | Normal | yes |
| Cyclin-dependent kinase 9 | Cdk9 | CG5179 | 34982 | Atrophy | no |
| Cyclin-dependent kinase 9 | Cdk9 | CG5179 | 35323 | Small ovaries | yes |
| Cyclin-dependent kinase 9 | Cdk9 | CG5179 | 41932 | Normal | yes |
| Cystathionine beta-synthase | Cbs | CG1753 | 36767 | Normal | yes |
| Cystathionine beta-synthase | Cbs | CG1753 | 41877 | Normal | yes |
| Cytochrome P450-4d8 | Cyp4d8 | CG4321 | 58019 | Atrophy | no |
| D-2-hydroxyglutaric acid dehydrogenase | D2hgdh | CG3835 | 53355 | Normal | yes |
| Darkener of apricot | Doa | CG42320 | 50903 | Normal | no |
| Dead box protein 80 | Dbp80 | CG17023 | 34682 | Normal | yes |
| deadhead | dhd | CG4193 | 41857 | Normal | yes |
| Death-associated inhibitor of apoptosis 1 / thread | Diap1 / th | CG12284 | 33597 | Normal | yes |
| decapentaplegic | dpp | CG9885 | 33618 | Normal | yes |
| decapentaplegic | dpp | CG9885 | 33767 | Normal | yes |
| decapentaplegic | dpp | CG9885 | 35214 | Normal | yes |
| Desaturase 1 | Desat1 | CG5887 | 35591 | Normal | yes |
| Desaturase 1 | Desat1 | CG5887 | 37512 | Normal | yes |
| diaphanous | dia | CG1768 | 33424 | Normal | no |
| diaphanous | dia | CG1768 | 35479 | Atrophy | no |
| Dicer-2 | Dcr2 | CG6493 | 33656 | Normal | yes |
| DISCO Interacting Protein 1 | DIP1 | CG17686 | 35226 | Normal | yes |
| DISCO Interacting Protein 1 | DIP1 | CG17686 | 35333 | Normal | yes |

| | | | | | |
|---|---|---|---|---|---|
| DISCO Interacting Protein 2 | DIP2 | CG7020 | 34918 | Normal | yes |
| DISCO Interacting Protein 2 | DIP2 | CG7020 | 42598 | Normal | yes |
| DnaJ-like-1 | Dnaj-1 | CG10578 | 32899 | Normal | yes |
| DnaJ-like-1 | Dnaj-1 | CG10578 | 32978 | Normal | yes |
| DnaJ-like-2 | Droj2 | CG8863 | 36089 | Normal | yes |
| Dodeca-satellite-binding protein 1 | Dp1 | CG5170 | 32872 | Normal | yes |
| domino | dom | CG9696 | 34827 | Atrophy | no |
| domino | dom | CG9696 | 38385 | Atrophy | no |
| domino | dom | CG9696 | 38941 | Normal | yes |
| domino | dom | CG9696 | 40914 | Atrophy | no |
| domino | dom | CG9696 | 41674 | Atrophy | no |
| dorsal | dl | CG6667 | 32934 | Normal | no |
| dorsal | dl | CG6667 | 36650 | Normal | no |
| dorsal | dl | CG6667 | 38905 | Normal | no |
| DP transcription factor | Dp | CG4654 | 33372 | Normal | yes |
| Dpt-YFP repressor by overexpression | Dyro | CG6175 | 62516 | Normal | yes |
| drosha | drosha | CG8730 | 33657 | Normal | nd |
| drosha | drosha | CG8730 | 35233 | Normal | yes |
| Dual oxidase | Duox | CG3131 | 32903 | Normal | yes |
| Dual oxidase | Duox | CG3131 | 33975 | Normal | yes |
| Dual oxidase | Duox | CG3131 | 38907 | Normal | yes |
| Dual oxidase | Duox | CG3131 | 38916 | Normal | yes |
| eIF4AIII | eIF4AIII | CG7483 | 32444 | Atrophy | no |
| eIF4AIII | eIF4AIII | CG7483 | 32907 | Atrophy | no |
| eIF4AIII | eIF4AIII | CG7483 | 38202 | Atrophy | no |
| Ejaculatory bulb protein III | PebIII | CG11390 | 55933 | Normal | yes |
| encore | enc | CG10847 | 42797 | Normal | yes |
| Enhancer of bithorax | E(bx) | CG32346 | 33658 | Normal | Subno |
| Enhancer of Polycomb | E(Pc) | CG7776 | 35271 | Atrophy | no |
| Enhancer of zeste | E(z) | CG6502 | 33659 | Normal | nd |
| Enhancer of zeste | E(z) | CG6502 | 36068 | Normal | no |
| enoki mushroom | enok | CG11290 | 40917 | Normal | yes |
| enoki mushroom | enok | CG11290 | 41664 | Normal | yes |
| enoki mushroom | enok | CG11290 | 42941 | Normal | yes |
| Ets at 97D | Ets97D | CG6338 | 35749 | Normal | yes |
| Ets at 97D | Ets97D | CG6338 | 36635 | Normal | yes |
| eukaryotic translation initiation factor 4E homologous protein | eIF4EHP | CG33100 | 36876 | Normal | nd |
| eukaryotic translation initiation factor 4E homologous protein | eIF4EHP | CG33100 | 43990 | Normal | yes |
| female sterile (1) Yb | fs(1)Yb | CG2706 | 35181 | Normal | yes |
| female sterile (1) Yb | fs(1)Yb | CG2706 | 35301 | Normal | yes |
| flower | fwe | CG6151 | 34157 | Normal | yes |
| flower | fwe | CG6151 | 43157 | Normal | yes |
| Fmr1 | Fmr1 | CG6203 | 35200 | Normal | yes |
| Frost | Fst | CG9434 | 33376 | Normal | yes |
| G9a | G9a | CG2995 | 34817 | Normal | yes |
| Gcn5 acetyltransferase / Pcaf | Gcn5 / Pcaf | CG4107 | 33981 | Normal | yes |
| Gcn5 acetyltransferase / Pcaf | Gcn5 / Pcaf | CG4107 | 35601 | Normal | no |
| Glutathione S transferase E1 | GstE1 | CG5164 | 36878 | Normal | yes |
| Glutathione S transferase E1 | GstE1 | CG5164 | 36878 | Normal | yes |
| Glutathione S transferase E6 | GstE6 | CG17530 | 65197 | Normal | yes |
| half pint | hfp | CG12085 | 34785 | Normal | yes |
| hangover | hang | CG32575 | 35674 | Normal | yes |
| hangover | hang | CG32575 | 41870 | Normal | no |
| haywire | hay | CG8019 | 53345 | Atrophy | no |
| Heat shock 70-kDa protein cognate 3 | Hsc70-3 | CG4147 | 32402 | Normal | yes |
| Heat shock factor | Hsf | CG5748 | 41581 | Normal | yes |
| Heat shock gene 67Ba | Hsp67Ba | CG4167 | 41962 | Normal | yes |
| Heat shock gene 67Ba | Hsp67Ba | CG4167 | 53007 | Normal | yes |
| Heat shock gene 67Bc | Hsp67Bc | CG4190 | 35452 | Normal | yes |
| Heat shock gene 67Bc | Hsp67Bc | CG4190 | 42607 | Normal | yes |
| Heat shock protein 22 | Hsp22 | CG4460 | 41709 | Normal | yes |
| Heat shock protein 22 | Hsp22 | CG4460 | 51397 | Normal | yes |
| Heat shock protein 23 | Hsp23 | CG4463 | 44029 | Normal | yes |
| Heat shock protein 26 | Hsp26 | CG4183 | 35408 | Normal | yes |
| Heat shock protein 26 | Hsp26 | CG4183 | 42610 | Normal | yes |
| Heat shock protein 27 | Hsp27 | CG4466 | 33007 | Normal | yes |

| | | | | | |
|---|---|---|---|---|---|
| Heat shock protein 27 | Hsp27 | CG4466 | 33922 | Normal | yes |
| Heat shock protein 60A | Hsp60A | CG12101 | 34729 | Normal | yes |
| Heat shock protein 60B | Hsp60B | CG2830 | 34729 | Normal | yes |
| Heat shock protein 67Bb | Hsp67Bb | CG4456 | 41709 | Normal | yes |
| Heat shock protein 67Bb | Hsp67Bb | CG4456 | 51397 | Normal | yes |
| Heat shock protein 68 | Hsp68 | CG5436 | 50637 | Normal | yes |
| Heat shock protein 70Aa | Hsp70Aa | CG31366 | 42639 | Normal | yes |
| Heat shock protein 70Ab | Hsp70Ab | CG18743 | 35663 | Normal | yes |
| Heat shock protein 70Ba | Hsp70Ba | CG31449 | 32997 | Normal | yes |
| Heat shock protein 70Ba | Hsp70Ba | CG31449 | 43289 | Normal | yes |
| Heat shock protein 70Ba | Hsp70Ba | CG31449 | 35672 | Normal | yes |
| Heat shock protein 70Bb | Hsp70Bb | CG31359 | 32997 | Normal | yes |
| Heat shock protein 70Bb | Hsp70Bb | CG31359 | 33948 | Normal | yes |
| Heat shock protein 70Bc | Hsp70Bc | CG6489 | 32997 | Normal | yes |
| Heat shock protein 70Bc | Hsp70Bc | CG6489 | 35697 | Normal | yes |
| Heat shock protein 70Bc | Hsp70Bc | CG6489 | 42626 | Normal | yes |
| Heat shock protein 83 | Hsp83 | CG1242 | 32996 | Normal | no |
| Heat shock protein 83 | Hsp83 | CG1242 | 33947 | Atrophy | no |
| Heat shock protein cognate 1 | Hsc70-1 | CG8937 | 34527 | Normal | yes |
| Heat shock protein cognate 2 | Hsc70-2 | CG7756 | 42014 | Normal | yes |
| Heat shock protein cognate 2 | Hsc70-2 | CG7756 | 44485 | Normal | yes |
| Heat shock protein cognate 2 | Hsc70-2 | CG7756 | 32997 | Normal | yes |
| Heat shock protein cognate 4 | Hsc70-4 | CG4264 | 34836 | Atrophy | no |
| Heat shock protein cognate 4 | Hsc70-4 | CG4264 | 35684 | Atrophy | no |
| Helicase at 25E | Hel25E | CG7269 | 33666 | Atrophy | no |
| hemipterous | hep | CG4353 | 35210 | Normal | no |
| hephaestus | heph | CG31000 | 35669 | Normal | no |
| hephaestus | heph | CG31000 | 55655 | Normal | yes |
| Heterochromatin Protein 1b | HP1b | CG7041 | 32401 | Normal | yes |
| Heterogeneous nuclear ribonucleoprotein at 27C | Hrb27C | CG10377 | 33716 | Normal | no |
| Heterogeneous nuclear ribonucleoprotein at 87F | Hrb87F | CG12749 | 52937 | Normal | yes |
| Heterogeneous nuclear ribonucleoprotein at 98DE | Hrb98DE | CG9983 | 32351 | Atrophy | no |
| Heterogeneous nuclear ribonucleoprotein K/bancal | HnRNP-K/bl | CG13425 | 42540 | Normal | yes |
| Host cell factor | Hcf | CG1710 | 32453 | Normal | yes |
| Host cell factor | Hcf | CG1710 | 36799 | Normal | yes |
| Hsp70Bbb | Hsp70Bbb | CG5834 | 32997 | Normal | yes |
| Hsp70Bbb | Hsp70Bbb | CG5834 | 33000 | Normal | yes |
| Hsp70Bbb | Hsp70Bbb | CG5834 | 33916 | Atrophy | no |
| HSPB1 associated protein 1 | HSPBAP1 | CG43320 | 34606 | Normal | yes |
| Imaginal disc growth factor 2 | Idgf2 | CG4475 | 55935 | Normal | yes |
| Imitation SWI | Iswi | CG8625 | 32845 | Normal | nd |
| Imitation SWI | Iswi | CG8625 | 51931 | Normal | yes |
| Insulin-like peptide 5 | Ilp5 | CG33273 | 33683 | Normal | yes |
| Inwardly rectifying potassium channel 1 | Irk1 | CG44159 | 42644 | Normal | yes |
| Isocitrate dehydrogenase | Idh | CG7176 | 41708 | Normal | yes |
| Jabba | Jabba | CG42351 | 36852 | Normal | no |
| janus A | janA | CG7933 | 41846 | Normal | yes |
| JIL-1 anchoring and stabilizing protein | Jasper | CG7946 | 55274 | Normal | nd |
| JIL-1 kinase | JIL-1 | CG6297 | 41592 | Normal | yes |
| JIL-1 kinase | JIL-1 | CG6297 | 42571 | ND | nd |
| JIL-1 kinase | JIL-1 | CG6297 | 55875 | Normal | yes |
| JIL-1 kinase | JIL-1 | CG6297 | 57293 | Normal | nd |
| jing interacting gene regulatory 1 | jigr1 | CG17383 | 58173 | Normal | yes |
| Jumonji domain containing 5 | JMJD5 | CG13902 | 33702 | Normal | yes |
| Jumonji, AT rich interactive domain 2 | jarid2 | CG3654 | 32891 | Normal | yes |
| Juvenile hormone epoxide hydrolase 3 | Jheh3 | CG15106 | 60021 | Normal | no |
| Kank | Kank | CG10249 | 33432 | Normal | yes |
| kayak | kay | CG33956 | 33379 | Normal | no |
| Keap1 | Keap1 | CG3962 | 40932 | Normal | no |
| kismet | kis | CG3696 | 34908 | Normal | yes |
| kismet | kis | CG3696 | 36597 | Normal | no |
| kismet | kis | CG3696 | 44542 | Normal | no |
| kohtalo | kto | CG8491 | 34588 | Normal | no |
| krimper | krimp | CG15707 | 35230 | Normal | no |
| krimper | krimp | CG15707 | 35231 | Normal | no |
| krimper | krimp | CG15707 | 37511 | Normal | no |

| | | | | | |
|---|---|---|---|---|---|
| Lamin | Lam | CG6944 | 36617 | Normal | yes |
| lethal (2) essential for life | l(2)efl | CG4533 | 41724 | Normal | yes |
| lethal (2) essential for life | l(2)efl | CG4533 | 51816 | Normal | yes |
| lethal (3) 72Ab | l(3)72Ab | CG5931 | 34024 | Atrophy | no |
| lethal (3) 72Ab | l(3)72Ab | CG5931 | 50716 | Atrophy | no |
| lethal (3) 80Fg | l(3)80Fg | CG40178 | 44578 | Normal | yes |
| Leucine zipper and EF-hand containing transmembrane protein 1 | Letm1 | CG4589 | 37502 | Atrophy | no |
| Ligase4 | lig4 | CG12176 | 51933 | Normal | yes |
| little imaginal discs | lid | CG9088 | 36652 | Normal | yes |
| little imaginal discs | lid | CG9088 | 35706 | Normal | yes |
| locomotion defects | loco | CG5248 | 32456 | Normal | yes |
| loki | lok | CG10895 | 35152 | Normal | no |
| longitudinals lacking | lola | CG12052 | 35721 | Normal | yes |
| loquacious | loqs | CG6866 | 32955 | Normal | yes |
| loquacious | loqs | CG6866 | 33427 | Normal | yes |
| loquacious | loqs | CG6866 | 34779 | Normal | yes |
| loquacious | loqs | CG6866 | 34780 | Normal | yes |
| loquacious | loqs | CG6866 | 34781 | Normal | yes |
| loquacious | loqs | CG6866 | 34782 | Normal | yes |
| loquacious | loqs | CG6866 | 34851 | Normal | yes |
| Lysine (K)-specific demethylase 2 | Kdm2 | CG11033 | 33699 | Normal | yes |
| Lysine (K)-specific demethylase 3 | Kdm3 | CG8165 | 32975 | Normal | no |
| Lysine (K)-specific demethylase 4A | Kdm4A | CG15835 | 34629 | Normal | yes |
| Lysine (K)-specific demethylase 4B | Kdm4B | CG33182 | 35676 | Normal | yes |
| mago nashi | mago | CG9401 | 35453 | Normal | yes |
| Major Facilitator Superfamily Transporter 17 | MFS17 | CG40263 | 44033 | Normal | yes |
| maleless | mle | CG11680 | 34864 | Normal | nd |
| maternal expression at 31B | Me31b | CG4916 | 33675 | Atrophy | no |
| maternal expression at 31B | Me31b | CG4916 | 38923 | Normal | yes |
| Max | Max | CG9648 | 40851 | Normal | yes |
| meiotic 41 | mei-41 | CG4252 | 35371 | Normal | yes |
| meiotic 41 | mei-41 | CG4252 | 41934 | Normal | yes |
| meiotic 9 | mei-9 | CG3697 | 55313 | Normal | yes |
| Mekk1 | Mekk1 | CG7717 | 35402 | Normal | yes |
| menage a trois | metro | CG30021 | 35810 | Normal | yes |
| Menin 1 | Mnn1 | CG13778 | 35150 | Atrophy | no |
| Menin 1 | Mnn1 | CG13778 | 51862 | Normal | yes |
| methuselah | mth | CG6936 | 36823 | Normal | yes |
| Methyltransferase 2 | Mt2 | CG10692 | 38224 | Normal | yes |
| Methyltransferase 2 | Mt2 | CG10692 | 42906 | Normal | yes |
| Mi-2 | Mi-2 | CG8103 | 35398 | Atrophy | no |
| Mi-2 | Mi-2 | CG8103 | 33419 | Atrophy | no |
| Mi-2 | Mi-2 | CG8103 | 51774 | Atrophy | no |
| moira | mor | CG18740 | 34919 | Normal | yes |
| moira | mor | CG18740 | 35630 | Normal | yes |
| moira | mor | CG18740 | 35662 | Normal | no |
| Mucin related 18B | Mur18B | CG7874 | 56957 | Normal | yes |
| mushroom-body expressed | mub | CG7437 | 34870 | Normal | yes |
| nejire | nej | CG15319 | 36682 | Normal | yes |
| nejire | nej | CG15319 | 37489 | Normal | no |
| Neurofibromin 1 | Nf1 | CG8318 | 53322 | Normal | yes |
| Nucleolar protein 66 | NO66 | CG2982 | 33596 | Normal | yes |
| Nucleosome remodeling factor - 38kD | Nurf-38 | CG4634 | 35444 | Normal | no |
| osa | osa | CG7467 | 35447 | Normal | yes |
| osa | osa | CG7467 | 38285 | Normal | yes |
| ovarian tumor | otu | CG12743 | 34065 | Atrophy | no |
| ovaries absent | ova | CG5694 | 36655 | Normal | yes |
| ovaries absent | ova | CG5694 | 62485 | Normal | yes |
| p23 | p23 | CG16817 | 41862 | Normal | yes |
| p24-related-2 | p24-2 | CG33105 | 40839 | Normal | yes |
| p38a MAP kinase | p38a | CG5475 | 34744 | Normal | yes |
| p38a MAP kinase | p38a | CG5475 | 35244 | Normal | yes |
| p38b MAP kinase | p38b | CG7393 | 35252 | Normal | yes |
| p53 | p53 | CG10895 | 36814 | Normal | yes |
| p53 | p53 | CG10895 | 41638 | Normal | yes |
| p53 | p53 | CG10895 | 41720 | Normal | yes |

| | | | | | |
|---|---|---|---|---|---|
| pacman | pcm | CG3291 | 34690 | Normal | yes |
| painless | pain | CG15860 | 51835 | Normal | yes |
| pancreatic eIF-2α kinase | PEK | CG2087 | 35162 | Normal | yes |
| pancreatic eIF-2α kinase | PEK | CG2087 | 42499 | Normal | yes |
| papi | papi | CG7082 | 34932 | Normal | yes |
| papi | papi | CG7082 | 35450 | Normal | yes |
| papi | papi | CG7082 | 37513 | Normal | yes |
| papi | papi | CG7082 | 38216 | Normal | yes |
| partner of drosha | pasha | CG1800 | 33972 | Normal | nd |
| peanuts | pea | CG8241 | 32838 | Atrophy | no |
| P-element induced wimpy testis | piwi | CG6122 | 33724 | Atrophy | no |
| P-element induced wimpy testis | piwi | CG6122 | 34866 | Atrophy | no |
| P-element induced wimpy testis | piwi | CG6122 | 37483 | Normal | no |
| P-element somatic inhibitor | Psi | CG8912 | 34825 | Normal | yes |
| Phosphoglycerate mutase 5 | Pgam5 | CG14816 | 33346 | Normal | yes |
| Phosphotidylinositol 3 kinase 59F | Pi3K59F | CG5373 | 33384 | Normal | yes |
| Phosphotidylinositol 3 kinase 59F | Pi3K59F | CG5373 | 36056 | Normal | yes |
| Poly-(ADP-ribose) polymerase | Parp | CG40411 | 35792 | Normal | yes |
| Polycomb | Pc | CG32443 | 33622 | Normal | yes |
| Polycomb | Pc | CG32443 | 33964 | Normal | yes |
| Polycomb | Pc | CG32443 | 36070 | Normal | yes |
| Polycomblike | Pcl | CG5109 | 33945 | Normal | yes |
| Polycomblike | Pcl | CG5109 | 33946 | Normal | yes |
| polyhomeotic proximal | ph-p | CG18412 | 33669 | Normal | yes |
| pontin | pont | CG4003 | 50972 | Atrophy | no |
| pre-mRNA processing factor 40 | Prp40 | CG3542 | 33711 | Normal | yes |
| pre-mRNA processing factor 6 | Prp6 | CG6841 | 51909 | Atrophy | no |
| Prip | Prip | CG7777 | 44464 | Normal | yes |
| Prip | Prip | CG7777 | 50695 | Normal | yes |
| protein partner of snf | pps | CG6525 | 38529 | Normal | yes |
| protein partner of snf | pps | CG6525 | 38912 | Normal | yes |
| puckered | puc | CG7850 | 53019 | Normal | yes |
| puckered | puc | CG7850 | 34392 | Normal | yes |
| puckered | puc | CG7850 | 36085 | Normal | no |
| puckered | puc | CG7850 | 44038 | Normal | yes |
| punt | put | CG7904 | 35195 | Normal | no |
| punt | put | CG7904 | 35701 | Normal | yes |
| punt | put | CG7904 | 39025 | Normal | yes |
| pyrexia | pyx | CG17142 | 51836 | Normal | yes |
| qin | qin | CG43726 | 37475 | Normal | yes |
| qin | qin | CG43726 | 41662 | Normal | nd |
| r2d2 | r2d2 | CG7138 | 34784 | Normal | yes |
| Rab5 | Rab5 | CG3664 | 34832 | Normal | no |
| Rab5 | Rab5 | CG3664 | 51847 | Normal | yes |
| raspberry | ras | CG1799 | 51717 | Small ovaries | yes |
| Ras-related protein interacting with calmodulin | Ric | CG8418 | 41819 | Normal | yes |
| Recombination repair protein 1 | Rrp1 | CG3178 | 35420 | Normal | yes |
| refractory to sigma P | ref(2)P | CG10360 | 33978 | Normal | yes |
| refractory to sigma P | ref(2)P | CG10360 | 36111 | Normal | yes |
| Regulator of telomere elongation helicase 1 | Rtel1 | CG4078 | 32973 | Normal | yes |
| Rev1 | Rev1 | CG12189 | 36654 | Normal | yes |
| rhino | rhi | CG10683 | 34071 | Normal | yes |
| rhino | rhi | CG10683 | 35171 | Normal | no |
| RhoGAP71E | RhoGAP71E | CG32149 | 32417 | Normal | yes |
| rho-type guanine exchange factor | rtGEF | CG10043 | 32947 | Normal | yes |
| Ribosomal protein L8 | Rpl8 | CG1263 | 50610 | Normal | yes |
| Rm62 | Rm62 | CG10279 | 34829 | Normal | yes |
| RNA polymerase II subunit Rpb4 | Rpb4 | CG43662 | 50905 | Atrophy | no |
| rotated abdomen | rt | CG6097 | 51805 | Normal | yes |
| Rox8 | Rox8 | CG5422 | 32472 | Normal | yes |
| Rpd3 | Rpd3 | CG7471 | 33725 | Normal | yes |
| Rpd3 | Rpd3 | CG7471 | 34846 | Normal | yes |
| Rrp6 | Rrp6 | CG7292 | 34809 | Letal | nd |
| Rrp6 | Rrp6 | CG7292 | 42064 | Normal | no |
| runt | run | CG1849 | 34707 | Normal | yes |
| sans fille | snf | CG4528 | 34593 | Normal | yes |

| | | | | | |
|---|---|---|---|---|---|
| sans fille | snf | CG4528 | 51459 | Atrophy | no |
| Scaffold attachment factor B | Saf-B | CG6995 | 51759 | Normal | no |
| Scamp | Scamp | CG9195 | 38277 | Normal | yes |
| scute | sc | CG3827 | 41594 | Normal | yes |
| Secreted protein, acidic, cysteine-rich | SPARC | CG6378 | 40885 | Normal | yes |
| Serine palmitoyltransferase subunit I | Spt-I | CG4016 | 55685 | Normal | yes |
| SET domain binding factor | Sbf | CG6939 | 32419 | Normal | yes |
| SET domain binding factor | Sbf | CG6939 | 44004 | Normal | yes |
| Sex lethal | Sxl | CG43770 | 34393 | Atrophy | no |
| Sex lethal | Sxl | CG43770 | 38195 | Atrophy | no |
| shibire | shi | CG18102 | 36921 | Normal | no |
| shotgun | shg | CG3722 | 32904 | Normal | yes |
| shotgun | shg | CG3722 | 38207 | Normal | no |
| shriveled | shv | CG4164 | 37507 | Normal | yes |
| shriveled | shv | CG4164 | 54797 | Normal | yes |
| shutdown | shu | CG4735 | 35454 | Normal | no |
| Sin3A | Sin3A | CG8815 | 32368 | Normal | yes |
| sisterless A | sisA | CG1641 | 55181 | Normal | yes |
| skuld | skd | CG9936 | 34630 | Normal | yes |
| slipper | slpr | CG2272 | 32948 | Normal | yes |
| slipper | slpr | CG2272 | 41605 | Normal | yes |
| small ribonucleoprotein particle U1 subunit 70K | snRNP-U1-70K | CG8749 | 33396 | Atrophy | no |
| small ribonucleoprotein particle U1 subunit C | snRNP-U1-C | CG5454 | 34822 | Atrophy | no |
| smaug | smg | CG5263 | 35477 | Normal | yes |
| Snf5-related 1 | Snr1 | CG1064 | 32372 | Normal | no |
| Son RNA binding protein | Son | CG8273 | 34805 | Normal | yes |
| Sorbitol dehydrogenase-2 | Sodh-2 | CG4649 | 53353 | Normal | Few yes |
| spaghetti squash | sqh | CG3595 | 32439 | ND | nd |
| spaghetti squash | sqh | CG3595 | 33892 | Normal | yes |
| spaghetti squash | sqh | CG3595 | 38222 | Normal | yes |
| Spf45 | Spf45 | CG17540 | 41954 | Normal | yes |
| spindle A | Spn-A | CG7948 | 38898 | Normal | yes |
| spindle A | Spn-A | CG7948 | 51936 | Normal | yes |
| spindle E | spn-E | CG3158 | 35303 | Normal | no |
| Splicing factor 2 | SF2 | CG6987 | 32367 | Normal | yes |
| Splicing factor 30 | Spf30 | CG17454 | 43199 | Atrophy | no |
| Splicing factor 3a subunit 1 | Sf3a1 | CG16941 | 34840 | Atrophy | no |
| Spt3 | Spt3 | CG3169 | 35148 | Normal | yes |
| Spt4 | Spt4 | CG12372 | 32896 | Atrophy | no |
| Spt5 | Spt5 | CG7626 | 34837 | Atrophy | no |
| Spt6 | Spt6 | CG12225 | 32373 | Atrophy | no |
| Spt7 | Spt7 | CG6506 | 42552 | Normal | yes |
| squid | sqd | CG16901 | 35627 | Normal | yes |
| squid | sqd | CG16901 | 53891 | Normal | yes |
| staufen | stau | CG5753 | 35690 | Normal | yes |
| staufen | stau | CG5753 | 43187 | Normal | no |
| stonewall | stwl | CG3836 | 35415 | Atrophy | no |
| Su(var)2-HP2 | Su(var)2-HP2 | CG12864 | 38255 | Atrophy | no |
| suppressor of Hairy wing | su(Hw) | CG8573 | 33906 | Normal | yes |
| suppressor of Hairy wing | su(Hw) | CG8573 | 34006 | Normal | yes |
| Suppressor of sable | su(sable) | CG6222 | 33982 | Normal | yes |
| Suppressor of Under-Replication | SuUR | CG7869 | 36893 | Normal | yes |
| Suppressor of variegation 205 | Su(var)205 | CG8409 | 33400 | Atrophy | no |
| Suppressor of variegation 205 | Su(var)205 | CG8409 | 36792 | Atrophy | no |
| Suppressor of variegation 2-10 | Su(var)2-10 | CG8068 | 32915 | Atrophy | no |
| Suppressor of variegation 2-10 | Su(var)2-10 | CG8068 | 32956 | Atrophy | no |
| Suppressor of variegation 2-10 | Su(var)2-10 | CG8068 | 58067 | Atrophy | no |
| Suppressor of variegation 3-3 | Su(var)3-3 | CG17149 | 32853 | Normal | yes |
| Suppressor of variegation 3-3 | Su(var)3-3 | CG17149 | 33726 | Normal | yes |
| Suppressor of variegation 3-3 | Su(var)3-3 | CG17149 | 36867 | Normal | yes |
| Suppressor of variegation 3-9 | Su(var)3-9 | CG43664 | 32914 | Atrophy | no |
| Suppressor of variegation 3-9 | Su(var)3-9 | CG43664 | 33401 | Atrophy | no |
| Suppressor of variegation 3-9 | Su(var)3-9 | CG43664 | 43661 | Atrophy | no |
| Suppressor of zeste 2 | Su(z)2 | CG3905 | 33403 | Normal | yes |
| survival motor neuron | Smn | CG16725 | 36621 | Normal | yes |

| | | | | | |
|---|---|---|---|---|---|
| tapas | tapas | CG8920 | 34738 | Normal | yes |
| TBP-associated factor 1 | Taf1 | CG17603 | 32421 | Atrophy | no |
| TBP-associated factor 1 | Taf1 | CG17603 | 35314 | Atrophy | no |
| TBP-associated factor 10 | Taf10 | CG2859 | 35239 | Normal | yes |
| TBP-associated factor 12 | Taf12 | CG17358 | 34852 | Normal | yes |
| TBP-associated factor 4 | Taf4 | CG5444 | 35427 | Normal | no |
| TBP-associated factor 4 | Taf4 | CG5444 | 50985 | Atrophy | no |
| TBP-associated factor 5 | Taf5 | CG7704 | 35367 | Atrophy | no |
| TBP-associated factor 7 | Taf7 | CG2670 | 55216 | Normal | yes |
| tejas | tej | CG8589 | 36879 | Normal | no |
| tejas | tej | CG8589 | 41929 | Normal | no |
| thickveins | tkv | CG14026 | 35166 | Normal | yes |
| thickveins | tkv | CG14026 | 35653 | Normal | yes |
| thickveins | tkv | CG14026 | 40937 | Atrophy | no |
| thickveins | tkv | CG14026 | 41904 | Normal | yes |
| Thioredoxin peroxidase 2 | Jafrac2 | CG1274 | 56043 | Normal | yes |
| Thor | Thor | CG8846 | 36667 | Normal | yes |
| Thor | Thor | CG8846 | 36815 | Normal | yes |
| transcriptional Adaptor 2a | Ada2a | CG43663 | 50905 | Atrophy | no |
| transcriptional Adaptor 2b | Ada2b | CG9638 | 35334 | Normal | yes |
| transcriptional Adaptor 3 | Ada3 | CG7098 | 32451 | Normal | yes |
| Transient receptor potential cation channel A1 | TrpA1 | CG5751 | 36780 | Normal | yes |
| trithorax | trx | CG8651 | 33703 | Normal | yes |
| tsunagi | tsu | CG8781 | 36585 | Atrophy | no |
| tudor | tud | CG9450 | 42800 | Normal | yes |
| Tudor domain containing 3 | Tdrd3 | CG13472 | 36819 | Normal | yes |
| Tudor staphylococcal nuclease | Tudor-SN | CG7008 | 34865 | Normal | nd |
| Turandot A | TotA | CG31509 | 53244 | Normal | yes |
| Turandot A | TotA | CG31509 | 55378 | Normal | yes |
| Turandot A | TotA | CG31509 | 58357 | Normal | yes |
| Turandot C | TotC | CG31508 | 51407 | Normal | yes |
| U2 small nuclear riboprotein auxiliary factor 38 | U2af38 | CG3582 | 50561 | Atrophy | no |
| U2 small nuclear riboprotein auxiliary factor 50 | U2af50 | CG9998 | 50521 | Normal | no |
| Uncoordinated 115a | Unc-115a | CG31352 | 40839 | Normal | yes |
| Uncoordinated 115b | Unc-115b | CG31332 | 40839 | Normal | yes |
| Upf1 | Upf1 | CG1559 | 43144 | Normal | no |
| Upf3 | Upf3 | CG11184 | 44565 | Normal | nd |
| upSET | upSET | CG9007 | 51447 | Normal | yes |
| Utx histone demethylase | Utx | CG5640 | 34076 | Normal | yes |
| vasa | vas | CG46283 | 32434 | Normal | no |
| vasa intronic gene | vig | CG4170 | 35183 | Normal | no |
| vasa intronic gene | vig | CG4170 | 35184 | Normal | yes |
| Victoria | Victoria | CG33117 | 55953 | Normal | yes |
| walrus | wal | CG8996 | 34915 | Normal | yes |
| windei | wde | CG12340 | 33339 | Normal | nd |
| XNP | XNP | CG4548 | 32894 | Normal | yes |
| β Hydroxy acid dehydrogenase 1 | Had1 | CG9914 | 62273 | Normal | yes |
| β-Mannosidase | β-Man | CG12582 | 53272 | Normal | yes |

**Extended Data Table 1. List of tested shRNA lines.**
