## Supplemental Table 2 for "The histone demethylase KDM3 prevents auto-immune piRNAs production in *Drosophila*"

| Library ID | Purpose | Genotype | Tissue | D. mel dm6 multimappers (depth) | Normalization factor (rpm) | D. mel dm6 clean 23-29nt unique mappers |
|---|---|---|---|---|---|---|
| GRH-103 | small seq | hfp GLKD | ovary | 8630134 | 0.116 | |
| GRH-104 | small seq | progeny of hfp GLKD | ovary | 12107109 | 0.083 | |
| GRH-105 | small seq | hfp control sisters | ovary | 10057434 | 0.099 | |
| GRH-106 | small seq | progeny of hfp control sisters | ovary | 16237313 | 0.062 | |
| GRH-111 | small seq | Kdm3 GLKD | ovary | 11418729 | 0.088 | |
| GRH-112 | small seq | Kdm3 control sisters | ovary | 14726077 | 0.068 | |
| ALBA-28 | small seq | Kdm3 GLKD | ovary | 20771697 | 0.048 | 1287136 |
| ALBA-29 | small seq | Kdm3 GLKD | ovary | 29582387 | 0.034 | 1996135 |
| ALBA-30 | small seq | Kdm3 GLKD | ovary | 25565165 | 0.039 | 637207 |
| ALBA-25 | small seq | Kdm3 control sisters | ovary | 24013259 | 0.042 | 1479172 |
| ALBA-26 | small seq | Kdm3 control sisters | ovary | 29159111 | 0.034 | 1754599 |
| ALBA-27 | small seq | Kdm3 control sisters | ovary | 23356824 | 0.043 | 1400470 |

| Library ID | Purpose | Genotype | Tissue | D. mel. dm6 reads | D. mel R6.36 all-genes reads |
|---|---|---|---|---|---|
| ALBA-1 | RNA seq | Kdm3 GLKD | ovary | 29044815 | 25268570 |
| ALBA-2 | RNA seq | Kdm3 GLKD | ovary | 41170212 | 34401515 |
| ALBA-3 | RNA seq | Kdm3 GLKD | ovary | 26531156 | 22946996 |
| ALBA-4 | RNA seq | Kdm3 control sisters | ovary | 33133242 | 27776026 |
| ALBA-5 | RNA seq | Kdm3 control sisters | ovary | 43201907 | 37041176 |
| ALBA-6 | RNA seq | Kdm3 control sisters | ovary | 38095053 | 31937164 |

| Library ID | Purpose | Genotype | Tissue | Antibody | D. mel. dm6 reads | D. mel. dm6 clean |
|---|---|---|---|---|---|---|
| ADQN-78 | ChIP seq | Kdm3 control sisters | ovary | no (INPUT for me2 and me3) | 38214851 | 37622419 |
| ADQN-79 | ChIP seq | Kdm3 GLKD | ovary | no (INPUT for me2 and me3) | 31085861 | 30561343 |
| ADQN-80 | ChIP seq | Kdm3 control sisters | ovary | αH3K9me2 | 17429811 | 15880104 |
| ADQN-81 | ChIP seq | Kdm3 control sisters | ovary | αH3K9me2 | 19637657 | 18181965 |
| ADQN-82 | ChIP seq | Kdm3 control sisters | ovary | αH3K9me2 | 21020633 | 19405338 |
| ADQN-83 | ChIP seq | Kdm3 GLKD | ovary | αH3K9me2 | 19386591 | 18602355 |
| ADQN-84 | ChIP seq | Kdm3 GLKD | ovary | αH3K9me2 | 25939857 | 24978229 |
| ADQN-85 | ChIP seq | Kdm3 GLKD | ovary | αH3K9me2 | 18977039 | 18211497 |
| ADQN-86 | ChIP seq | Kdm3 control sisters | ovary | αH3K9me3 | 17114226 | 15827951 |
| ADQN-87 | ChIP seq | Kdm3 control sisters | ovary | αH3K9me3 | 19588057 | 18078106 |
| ADQN-88 | ChIP seq | Kdm3 control sisters | ovary | αH3K9me3 | 23214725 | 21711093 |
| ADQN-89 | ChIP seq | Kdm3 GLKD | ovary | αH3K9me3 | 22208263 | 20666374 |
| ADQN-90 | ChIP seq | Kdm3 GLKD | ovary | αH3K9me3 | 24629347 | 23093308 |
| ADQN-91 | ChIP seq | Kdm3 GLKD | ovary | αH3K9me3 | 23634231 | 22259230 |
| S1 | ChIP seq | Kdm3 control sisters | ovary | no (INPUT for Rhino) | 21871351 | 21491195 |
| S2 | ChIP seq | Kdm3 control sisters | ovary | αRhino | 21389050 | 20979220 |
| S3 | ChIP seq | Kdm3 control sisters | ovary | αRhino | 20097195 | 19719881 |
| S4 | ChIP seq | Kdm3 GLKD | ovary | no (INPUT for Rhino) | 22166658 | 21797391 |
| S5 | ChIP seq | Kdm3 GLKD | ovary | αRhino | 19800458 | 19457139 |
| S6 | ChIP seq | Kdm3 GLKD | ovary | αRhino | 18726314 | 18378218 |

**Extended Data Table 2. Summary of small RNAseq, RNAseq and ChIPseq data.**
