## Supplemental Table 3 for "The histone demethylase KDM3 prevents auto-immune piRNAs production in *Drosophila*"

| Generation | Repression (%) | n |
|---|---|---|
| 3 | 100 | 46 |
| 4 | 100 | 50 |
| 8 | 100 | 44 |
| 20 | 100 | 72 |

**Extended Data Table 3. Stability of *BX2* conversion through subsequent generations.**
