## Supplemental Table 4 for "The histone demethylase KDM3 prevents auto-immune piRNAs production in *Drosophila*"

| Region | Size | Rank | GeneContent | ID | MeanControl (rpm) | MeanMutant (rpm) | SigmaControl | SigmaMutant | BaseMean | FC | log2FC | log2control | log2mutant |
|---|---|---|---|---|---|---|---|---|---|---|---|---|---|
| chr2L:21766000-21948000 | 182000 | R1 | tsh | R1_tsh | 17.360 | 531.233 | 2.922 | 22.379 | 274.297 | 30.600 | 4.935 | 4.118 | 9.053 |
| chrX:4372000-4532000 | 160000 | R2 | bi | R2_bi | 7.060 | 672.767 | 3.505 | 46.215 | 339.914 | 95.294 | 6.574 | 2.820 | 9.394 |
| chrX:16101000-16223000 | 122000 | R3 | disco-r/disco | R3_disco-r/disco | 7.591 | 272.479 | 1.746 | 25.798 | 140.035 | 35.897 | 5.166 | 2.924 | 8.090 |
| chr2R:6503000-6613000 | 110000 | R4 | jing | R4_jing | 7.717 | 566.480 | 2.784 | 24.736 | 287.099 | 73.404 | 6.198 | 2.948 | 9.146 |
| chrX:7160000-7236000 | 76000 | R5 | CR44357 | R5_CR44357 | 0.619 | 61.925 | 0.281 | 0.192 | 31.272 | 100.023 | 6.644 | -0.692 | 5.952 |
| chrX:8742000-8812000 | 70000 | R6 | Lim1 | R6_Lim1 | 2.224 | 87.556 | 0.305 | 9.334 | 44.890 | 39.376 | 5.299 | 1.153 | 6.452 |
| chr3R:4822000-4888000 | 66000 | R7 | opa/laf | R7_opa/laf | 3.589 | 420.565 | 2.122 | 15.841 | 212.077 | 117.173 | 6.872 | 1.844 | 8.716 |
| chr3L:22882000-22945000 | 63000 | R8 | BoYb | R8_BoYb | 7.472 | 186.016 | 0.985 | 8.605 | 96.744 | 24.896 | 4.638 | 2.901 | 7.539 |
| chrX:17767000-17820000 | 53000 | R9 | unc4/OdsH | R9_unc4/OdsH | 2.347 | 75.834 | 0.800 | 8.779 | 39.091 | 32.307 | 5.014 | 1.231 | 6.245 |
| chr4:491000-538000 | 47000 | R10 | zhf2 | R10_zhf2 | 1.832 | 163.540 | 0.405 | 3.116 | 82.686 | 89.281 | 6.480 | 0.873 | 7.353 |
| chr3L:23443000-23487000 | 44000 | R-1 | intergenic | R-1_intergenic | 211.174 | 15.835 | 7.082 | 1.613 | 113.505 | 0.075 | -3.737 | 7.722 | 3.985 |
| chr2R:5718000-5753000 | 35000 | R11 | ap | R11_ap | 2.451 | 90.000 | 1.027 | 4.701 | 46.225 | 36.726 | 5.199 | 1.293 | 6.492 |
| chr3R:4480000-4515000 | 35000 | R12 | Fip1 | R12_Fip1 | 2.116 | 67.709 | 0.679 | 2.177 | 34.912 | 32.003 | 5.000 | 1.081 | 6.081 |
| chrX:518000-552000 | 34000 | R13 | Appl | R13_Appl | 1.246 | 39.994 | 0.111 | 9.320 | 20.620 | 32.110 | 5.005 | 0.317 | 5.322 |
| chr3L:22828000-22862000 | 34000 | R14 | jim | R14_jim | 3.756 | 69.363 | 0.491 | 5.799 | 36.560 | 18.466 | 4.207 | 1.909 | 6.116 |
| chr2L:21991000-22019000 | 28000 | R15 | CG31693 | R15_CG31693 | 0.598 | 27.039 | 0.203 | 1.313 | 13.818 | 45.251 | 5.500 | -0.743 | 4.757 |
| chr2R:8636000-8664000 | 28000 | R16 | ptc | R16_ptc | 1.283 | 24.849 | 0.142 | 6.610 | 13.066 | 19.373 | 4.276 | 0.359 | 4.635 |
| chrX:2186000-2214000 | 28000 | R17 | CG14053 | R17_CG14053 | 4.250 | 49.671 | 0.257 | 5.096 | 26.961 | 11.688 | 3.547 | 2.087 | 5.634 |
| chrX:16231000-16258000 | 27000 | R18 | snRNA U5 | R18_snRNA U5 | 0.980 | 25.068 | 0.162 | 1.000 | 13.024 | 25.583 | 4.677 | -0.029 | 4.648 |
| chr3R:4341000-4366000 | 25000 | R19 | CG1090 | R19_CG1090 | 0.433 | 27.795 | 0.142 | 2.544 | 14.114 | 64.185 | 6.004 | -1.207 | 4.797 |
| chr2L:22029000-22054000 | 25000 | R20 | CG31601 | R20_CG31601 | 0.372 | 16.198 | 0.144 | 0.830 | 8.285 | 43.524 | 5.444 | -1.426 | 4.018 |
| chr3R:4914000-4938000 | 24000 | R21 | Cdep | R21_Cdep | 1.618 | 18.290 | 0.031 | 1.442 | 9.954 | 11.303 | 3.499 | 0.694 | 4.193 |
| chrX:9283000-9306000 | 23000 | R22 | lz | R22_lz | 0.844 | 47.403 | 0.072 | 3.668 | 24.123 | 56.194 | 5.812 | -0.245 | 5.567 |
| chr2R:13019000-13042000 | 23000 | R23 | intergenic | R23_intergenic | 1.693 | 21.990 | 0.159 | 2.318 | 11.841 | 12.990 | 3.699 | 0.759 | 4.459 |
| chrX:8627000-8649000 | 22000 | R24 | oc | R24_oc | 0.151 | 13.269 | 0.132 | 3.427 | 6.710 | 87.653 | 6.454 | -2.724 | 3.730 |
| chr3R:4278000-4300000 | 22000 | R25 | cpx | R25_cpx | 2.647 | 28.422 | 0.306 | 3.258 | 15.535 | 10.736 | 3.424 | 1.405 | 4.829 |
| chr3L:22750000-22772000 | 22000 | R26 | SpoCk | R26_SpoCk | 2.919 | 28.406 | 0.838 | 1.549 | 15.663 | 9.730 | 3.282 | 1.546 | 4.828 |
| chr3R:30249000-30268000 | 19000 | R27 | CG18404 | R27_CG18404 | 2.644 | 27.128 | 0.599 | 3.106 | 14.886 | 10.261 | 3.359 | 1.403 | 4.762 |
| chr3R:12357000-12376000 | 19000 | R28 | GstD9 | R28_GstD9 | 16.945 | 131.013 | 0.865 | 18.087 | 73.979 | 7.732 | 2.951 | 4.083 | 7.034 |
| chrX:4599000-4617000 | 18000 | R29 | CR32773 | R29_CR32773 | 0.164 | 6.158 | 0.065 | 1.160 | 3.161 | 37.438 | 5.226 | -2.604 | 2.622 |
| chr3L:371000-389000 | 18000 | R30 | trh | R30_trh | 0.305 | 11.358 | 0.154 | 2.852 | 5.831 | 37.247 | 5.219 | -1.713 | 3.506 |
| chrX:22832000-22850000 | 18000 | R31 | fog | R31_fog | 0.796 | 12.570 | 0.200 | 1.801 | 6.683 | 15.794 | 3.981 | -0.329 | 3.652 |
| chr3L:1448000-1466000 | 18000 | R32 | rho | R32_rho | 0.518 | 7.754 | 0.248 | 0.653 | 4.136 | 14.958 | 3.903 | -0.948 | 2.955 |
| chr3L:21768000-21786000 | 18000 | R33 | Syn1 | R33_Syn1 | 0.875 | 12.265 | 0.118 | 1.426 | 6.570 | 14.012 | 3.809 | -0.192 | 3.617 |
| chr2L:19013000-19030000 | 17000 | R34 | CR43700 | R34_CR43700 | 0.209 | 9.235 | 0.004 | 0.335 | 4.722 | 44.114 | 5.463 | -2.256 | 3.207 |
| chrX:6009000-6026000 | 17000 | R35 | mab21 | R35_mab21 | 0.219 | 8.712 | 0.198 | 0.790 | 4.466 | 39.749 | 5.313 | -2.190 | 3.123 |
| chr3R:4241000-4258000 | 17000 | R36 | TwdlG | R36_TwdlG | 1.587 | 50.252 | 0.257 | 0.868 | 25.919 | 31.664 | 4.985 | 0.666 | 5.651 |
| chr3L:1375000-1392000 | 17000 | R37 | Ptp61F | R37_Ptp61F | 0.870 | 13.554 | 0.542 | 0.807 | 7.212 | 15.575 | 3.961 | -0.200 | 3.761 |
| chr3L:19481000-19498000 | 17000 | R38 | CG9449 | R38_CG9449 | 1.204 | 15.196 | 0.183 | 2.740 | 8.200 | 12.618 | 3.657 | 0.268 | 3.926 |
| chr2L:20231000-20247000 | 16000 | R39 | intergenic | R39_intergenic | 0.281 | 13.292 | 0.087 | 1.798 | 6.787 | 47.341 | 5.565 | -1.832 | 3.733 |
| chrX:568000-584000 | 16000 | R40 | Appl | R40_Appl | 1.636 | 16.405 | 0.339 | 2.409 | 9.020 | 10.030 | 3.326 | 0.710 | 4.036 |
| chr3L:64000-79000 | 15000 | R-2 | Lsp1gamma | R-2_Lsp1gamma | 34.777 | 0.954 | 7.728 | 0.166 | 17.866 | 0.027 | -5.188 | 5.120 | -0.068 |
| chr3R:6662000-6677000 | 15000 | R41 | lab | R41_lab | 0.325 | 58.850 | 0.343 | 12.379 | 29.587 | 181.346 | 7.503 | -1.624 | 5.879 |
| chr2L:20257000-20272000 | 15000 | R42 | CG17570 | R42_CG17570 | 0.091 | 4.870 | 0.086 | 0.609 | 2.481 | 53.297 | 5.736 | -3.452 | 2.284 |
| chrX:766000-781000 | 15000 | R43 | fz3 | R43_fz3 | 0.619 | 22.312 | 0.068 | 0.327 | 11.466 | 36.019 | 5.171 | -0.691 | 4.480 |
| chr3R:3897000-3911000 | 14000 | R-3 | intergenic | R-3_intergenic | 592.201 | 46.029 | 20.520 | 6.111 | 319.115 | 0.078 | -3.685 | 9.210 | 5.524 |
| chr3L:22078000-22092000 | 14000 | R44 | msopa | R44_msopa | 0.144 | 8.107 | 0.020 | 0.599 | 4.126 | 56.273 | 5.814 | -2.795 | 3.019 |
| chr2L:19071000-19085000 | 14000 | R45 | Lim3 | R45_Lim3 | 0.173 | 7.186 | 0.119 | 0.730 | 3.680 | 41.428 | 5.373 | -2.527 | 2.845 |
| chr3L:21202000-21216000 | 14000 | R46 | CG10508 | R46_CG10508 | 0.616 | 15.844 | 0.169 | 0.935 | 8.230 | 25.707 | 4.684 | -0.698 | 3.986 |
| chr4:402000-416000 | 14000 | R47 | intergenic | R47_intergenic | 0.565 | 14.038 | 0.088 | 1.930 | 7.302 | 24.854 | 4.635 | -0.824 | 3.811 |
| chr3R:18911000-18925000 | 14000 | R48 | Xrp1 | R48_Xrp1 | 3.600 | 33.096 | 0.662 | 3.134 | 18.348 | 9.193 | 3.201 | 1.848 | 5.049 |
| chr3R:3376000-3389000 | 13000 | R-4 | intergenic | R-4_intergenic | 232.767 | 34.188 | 6.463 | 2.647 | 133.477 | 0.147 | -2.767 | 7.863 | 5.095 |
| chrX:8608000-8621000 | 13000 | R49 | intergenic | R49_intergenic | 0.124 | 11.910 | 0.109 | 3.621 | 6.017 | 95.959 | 6.584 | -3.010 | 3.574 |
| chr3R:4894000-4907000 | 13000 | R50 | CR45580 | R50_CR45580 | 0.717 | 58.559 | 0.394 | 2.617 | 29.638 | 81.686 | 6.352 | -0.480 | 5.872 |
| chr2R:8784000-8797000 | 13000 | R51 | intergenic | R51_intergenic | 0.276 | 7.065 | 0.168 | 2.055 | 3.670 | 25.643 | 4.680 | -1.860 | 2.821 |
| chr3R:4449000-4462000 | 13000 | R52 | CG31522 | R52_CG31522 | 0.370 | 8.019 | 0.055 | 1.239 | 4.195 | 21.665 | 4.437 | -1.434 | 3.003 |
| chrX:21504000-21517000 | 13000 | R53 | CR45082 | R53_CR45082 | 0.457 | 6.950 | 0.176 | 1.016 | 3.703 | 15.220 | 3.928 | -1.131 | 2.797 |
| chr2L:22059000-22071000 | 12000 | R54 | CG42597 | R54_CG42597 | 0.131 | 4.354 | 0.044 | 0.484 | 2.242 | 33.347 | 5.059 | -2.937 | 2.122 |
| chr3L:16762000-16774000 | 12000 | R55 | Nrt | R55_Nrt | 0.157 | 4.823 | 0.126 | 0.894 | 2.490 | 30.689 | 4.940 | -2.670 | 2.270 |
| chrX:22719000-22731000 | 12000 | R56 | CR44997 | R56_CR44997 | 1.222 | 12.244 | 0.297 | 0.434 | 6.733 | 10.017 | 3.324 | 0.290 | 3.614 |
| chr3R:3989000-4000000 | 11000 | R-5 | intergenic | R-5_intergenic | 87.920 | 5.194 | 4.903 | 0.299 | 46.557 | 0.059 | -4.081 | 6.458 | 2.377 |
| chr3R:6964000-6975000 | 11000 | R57 | Antp | R57_Antp | 0.158 | 15.627 | 0.154 | 0.487 | 7.893 | 98.652 | 6.624 | -2.658 | 3.966 |
| chr2L:21200000-21211000 | 11000 | R58 | clumsy | R58_clumsy | 0.147 | 6.656 | 0.038 | 1.071 | 3.402 | 45.311 | 5.502 | -2.767 | 2.735 |
| chrX:20629000-20640000 | 11000 | R59 | intergenic | R59_intergenic | 0.167 | 6.915 | 0.039 | 0.234 | 3.541 | 41.427 | 5.372 | -2.583 | 2.790 |
| chr3L:24555000-24566000 | 11000 | R-6 | intergenic | R-6_intergenic | 507.427 | 16.506 | 20.106 | 0.846 | 261.967 | 0.033 | -4.942 | 8.987 | 4.045 |
| chr2L:21971000-21982000 | 11000 | R60 | CG2528 | R60_CG2528 | 0.254 | 7.476 | 0.185 | 0.613 | 3.865 | 29.456 | 4.880 | -1.978 | 2.902 |
| chr3L:21251000-21262000 | 11000 | R61 | Eip78C | R61_Eip78C | 1.270 | 26.724 | 0.284 | 3.449 | 13.997 | 21.041 | 4.395 | 0.345 | 4.740 |
| chrX:18617000-18628000 | 11000 | R62 | wgn | R62_wgn | 0.324 | 5.453 | 0.102 | 0.292 | 2.889 | 16.804 | 4.071 | -1.624 | 2.447 |
| chr2L:14477000-14488000 | 11000 | R63 | CR44731 | R63_CR44731 | 0.829 | 7.637 | 0.288 | 1.087 | 4.233 | 9.214 | 3.204 | -0.271 | 2.933 |
| chr2L:492000-503000 | 11000 | R64 | ush | R64_ush | 0.689 | 5.352 | 0.126 | 0.910 | 3.021 | 7.765 | 2.957 | -0.537 | 2.420 |
| chrX:8728000-8738000 | 10000 | R65 | intergenic | R65_intergenic | 0.054 | 6.620 | 0.028 | 1.032 | 3.337 | 122.915 | 6.942 | -4.215 | 2.727 |
| chr3L:22274000-22284000 | 10000 | R66 | Sprk79D | R66_Sprk79D | 0.334 | 17.260 | 0.077 | 1.947 | 8.797 | 51.699 | 5.692 | -1.583 | 4.109 |
| chrX:7141000-7151000 | 10000 | R67 | CR32730 | R67_CR32730 | 0.140 | 5.245 | 0.087 | 0.214 | 2.692 | 37.580 | 5.232 | -2.841 | 2.391 |
| chr2R:11391000-11401000 | 10000 | R68 | sprt | R68_sprt | 0.111 | 3.857 | 0.085 | 0.836 | 1.984 | 34.864 | 5.124 | -3.176 | 1.947 |
| chrX:22513000-22523000 | 10000 | R69 | CG17600 | R69_CG17600 | 0.318 | 7.041 | 0.141 | 0.554 | 3.680 | 22.121 | 4.467 | -1.652 | 2.816 |
| chr3R:3878000-3887000 | 9000 | R-7 | intergenic | R-7_intergenic | 141.220 | 15.192 | 6.881 | 1.030 | 78.206 | 0.108 | -3.217 | 7.142 | 3.925 |
| chrX:5601000-5610000 | 9000 | R70 | Vsx1 | R70_Vsx1 | 0.000 | 3.253 | 0.000 | 1.093 | 1.627 | nc | nc | nc | 1.702 |
| chr3L:278000-287000 | 9000 | R71 | RhoGEF3 | R71_RhoGEF3 | 0.039 | 3.064 | 0.042 | 0.196 | 1.552 | 78.172 | 6.289 | -4.673 | 1.615 |
| chr3L:9012000-9021000 | 9000 | R72 | Doc2 | R72_Doc2 | 0.043 | 3.303 | 0.074 | 0.695 | 1.673 | 77.150 | 6.270 | -4.546 | 1.724 |
| chrX:22786000-22795000 | 9000 | R73 | CR44997 | R73_CR44997 | 0.167 | 8.645 | 0.192 | 0.594 | 4.406 | 51.643 | 5.691 | -2.579 | 3.112 |
| chrX:22693000-22702000 | 9000 | R74 | CR44997 | R74_CR44997 | 0.207 | 4.060 | 0.072 | 0.888 | 2.134 | 19.587 | 4.292 | -2.270 | 2.022 |
| chr2L:20128000-20137000 | 9000 | R75 | CG10651 | R75_CG10651 | 0.269 | 4.330 | 0.067 | 0.170 | 2.299 | 16.098 | 4.009 | -1.895 | 2.114 |

| | | | | | | | | | | | | | |
|---|---|---|---|---|---|---|---|---|---|---|---|---|---|
| chr2L:16477000-16485000 | 8000 | R76 | dac | R76_dac | 0.040 | 3.397 | 0.043 | 0.774 | 1.718 | 84.975 | 6.409 | -4.645 | 1.764 |
| chrX:17755000-17763000 | 8000 | R77 | intergenic | R77_intergenic | 0.062 | 4.815 | 0.035 | 0.743 | 2.439 | 77.108 | 6.269 | -4.001 | 2.268 |
| chrX:21482000-21490000 | 8000 | R78 | CR45511 | R78_CR45511 | 0.051 | 3.597 | 0.044 | 0.768 | 1.824 | 71.049 | 6.151 | -4.304 | 1.847 |
| chr2R:25084000-25092000 | 8000 | R79 | lov | R79_lov | 0.042 | 2.903 | 0.072 | 0.269 | 1.472 | 69.712 | 6.123 | -4.586 | 1.538 |
| chr2R:4874000-4882000 | 8000 | R-8 | intergenic | R-8_intergenic | 20.197 | 2.840 | 2.836 | 0.807 | 11.518 | 0.141 | -2.830 | 4.336 | 1.506 |
| chr2L:21185000-21193000 | 8000 | R80 | Ret | R80_Ret | 0.060 | 3.517 | 0.070 | 0.664 | 1.788 | 58.996 | 5.883 | -4.068 | 1.814 |
| chr3R:31158000-31166000 | 8000 | R81 | stops | R81_stops | 0.037 | 1.995 | 0.035 | 0.187 | 1.016 | 54.286 | 5.762 | -4.766 | 0.996 |
| chr3L:21357000-21365000 | 8000 | R82 | CG32440 | R82_CG32440 | 0.269 | 11.341 | 0.088 | 0.912 | 5.805 | 42.226 | 5.400 | -1.897 | 3.503 |
| chrX:356000-364000 | 8000 | R83 | y | R83_y | 0.088 | 2.713 | 0.047 | 0.293 | 1.400 | 30.910 | 4.950 | -3.510 | 1.440 |
| chrX:1381000-1389000 | 8000 | R84 | CG32813 | R84_CG32813 | 0.330 | 7.503 | 0.076 | 1.124 | 3.917 | 22.725 | 4.506 | -1.599 | 2.907 |
| chr2R:6058000-6066000 | 8000 | R85 | CCHa2 | R85_CCHa2 | 0.149 | 2.895 | 0.139 | 0.090 | 1.522 | 19.490 | 4.285 | -2.751 | 1.534 |
| chr2R:24950000-24958000 | 8000 | R86 | CG12851 | R86_CG12851 | 0.122 | 2.260 | 0.117 | 0.438 | 1.191 | 18.466 | 4.207 | -3.031 | 1.176 |
| chr2R:8327000-8335000 | 8000 | R87 | pdm3 | R87_pdm3 | 0.170 | 2.786 | 0.003 | 0.183 | 1.478 | 16.410 | 4.037 | -2.558 | 1.478 |
| chr2R:5790000-5798000 | 8000 | R88 | Or42a | R88_Or42a | 0.287 | 4.200 | 0.153 | 0.295 | 2.244 | 14.622 | 3.870 | -1.800 | 2.070 |
| chr2L:21709000-21717000 | 8000 | R89 | nolo | R89_nolo | 0.362 | 5.244 | 0.105 | 0.679 | 2.803 | 14.505 | 3.858 | -1.468 | 2.391 |
| chrX:868000-876000 | 8000 | R90 | CG11664 | R90_CG11664 | 0.305 | 4.229 | 0.068 | 1.725 | 2.267 | 13.871 | 3.794 | -1.714 | 2.080 |
| chr4:385000-393000 | 8000 | R91 | dati | R91_dati | 0.394 | 4.506 | 0.309 | 0.401 | 2.450 | 11.437 | 3.516 | -1.344 | 2.172 |
| chr2R:23622000-23630000 | 8000 | R92 | intergenic | R92_intergenic | 0.202 | 2.212 | 0.096 | 0.598 | 1.207 | 10.972 | 3.456 | -2.310 | 1.145 |
| chrX:2790000-2797000 | 7000 | R100 | w(BX2) | R100_w(BX2) | 6.426 | 283.014 | 0.852 | 44.673 | 144.720 | 44.040 | 5.461 | 2.684 | 8.145 |
| chrX:10619000-10626000 | 7000 | R101 | Rhab9Db | R101_Rhab9Db | 0.116 | 4.942 | 0.029 | 0.885 | 2.529 | 42.492 | 5.409 | -3.104 | 2.305 |
| chr3R:4778000-4785000 | 7000 | R102 | CG17387 | R102_CG17387 | 0.088 | 3.674 | 0.047 | 0.498 | 1.881 | 41.866 | 5.388 | -3.510 | 1.877 |
| chr2L:22075000-22082000 | 7000 | R103 | ttm3 | R103_ttm3 | 0.090 | 3.283 | 0.043 | 0.297 | 1.687 | 36.394 | 5.186 | -3.471 | 1.715 |
| chr3L:21237000-21244000 | 7000 | R104 | Eip78C | R104_Eip78C | 0.345 | 9.711 | 0.151 | 1.556 | 5.028 | 28.189 | 4.817 | -1.537 | 3.280 |
| chr3R:16217000-16224000 | 7000 | R105 | CR45643 | R105_CR45643 | 0.085 | 2.292 | 0.086 | 0.281 | 1.189 | 27.018 | 4.756 | -3.559 | 1.197 |
| chr3R:4520000-4527000 | 7000 | R106 | CG34357 | R106_CG34357 | 0.255 | 5.472 | 0.076 | 0.844 | 2.863 | 21.451 | 4.423 | -1.971 | 2.452 |
| chr2R:11486000-11493000 | 7000 | R107 | inv | R107_inv | 0.161 | 2.833 | 0.053 | 0.477 | 1.497 | 17.618 | 4.139 | -2.637 | 1.502 |
| chrX:1534000-1541000 | 7000 | R108 | Mur2B | R108_Mur2B | 0.294 | 5.059 | 0.090 | 1.446 | 2.676 | 17.194 | 4.104 | -1.765 | 2.339 |
| chr2L:435000-442000 | 7000 | R109 | ex | R109_ex | 0.207 | 2.872 | 0.034 | 0.481 | 1.539 | 13.905 | 3.798 | -2.276 | 1.522 |
| chr2L:9774000-9781000 | 7000 | R110 | ppk | R110_ppk | 0.351 | 4.620 | 0.241 | 0.820 | 2.486 | 13.164 | 3.718 | -1.510 | 2.208 |
| chr2R:7246000-7253000 | 7000 | R111 | Gadd45a | R111_Gadd45a | 4.787 | 51.388 | 0.557 | 7.193 | 28.087 | 10.736 | 3.424 | 2.259 | 5.683 |
| chr2R:24891000-24898000 | 7000 | R112 | Ance-5 | R112_Ance-5 | 0.344 | 3.687 | 0.100 | 0.997 | 2.016 | 10.717 | 3.422 | -1.539 | 1.883 |
| chr2L:21607000-21614000 | 7000 | R113 | intergenic | R113_intergenic | 0.875 | 8.135 | 0.036 | 0.367 | 4.505 | 9.302 | 3.218 | -0.193 | 3.024 |
| chr3L:14614000-14621000 | 7000 | R114 | shd | R114_shd | 0.462 | 3.484 | 0.120 | 1.119 | 1.973 | 7.543 | 2.915 | -1.114 | 1.801 |
| chr3L:21269000-21276000 | 7000 | R115 | AcCoAS | R115_AcCoAS | 3.326 | 6.537 | 0.512 | 0.425 | 4.931 | 1.965 | 0.975 | 1.734 | 2.709 |
| chr3R:16138000-16145000 | 7000 | R-9 | Sb | R-9_Sb | 10.060 | 0.098 | 2.379 | 0.063 | 5.079 | 0.010 | -6.675 | 3.331 | -3.344 |
| chr2L:19515000-19522000 | 7000 | R93 | CG10132 | R93_CG10132 | 0.011 | 2.859 | 0.020 | 0.767 | 1.435 | 250.117 | 7.966 | -6.451 | 1.516 |
| chr3R:31263000-31270000 | 7000 | R94 | ppk24 | R94_ppk24 | 0.023 | 3.944 | 0.040 | 0.271 | 1.983 | 172.506 | 7.431 | -5.451 | 1.980 |
| chrX:7242000-7249000 | 7000 | R95 | CR44357 | R95_CR44357 | 0.023 | 2.781 | 0.040 | 0.330 | 1.402 | 121.619 | 6.926 | -5.451 | 1.475 |
| chrX:5533000-5540000 | 7000 | R96 | Vsx2 | R96_Vsx2 | 0.040 | 2.747 | 0.005 | 0.476 | 1.393 | 69.399 | 6.117 | -4.659 | 1.458 |
| chr2R:14781000-14788000 | 7000 | R97 | kn | R97_kn | 0.068 | 3.334 | 0.052 | 0.329 | 1.701 | 48.943 | 5.613 | -3.876 | 1.737 |
| chr3R:5437000-5444000 | 7000 | R98 | CG14669 | R98_CG14669 | 0.077 | 3.690 | 0.032 | 0.955 | 1.883 | 48.095 | 5.588 | -3.704 | 1.884 |
| chrX:9310000-9317000 | 7000 | R99 | CR44534 | R99_CR44534 | 0.099 | 4.621 | 0.089 | 0.235 | 2.360 | 46.769 | 5.547 | -3.339 | 2.208 |
| chr3R:3972000-3978000 | 6000 | R-10 | intergenic | R-10_intergenic | 10.240 | 1.364 | 0.524 | 0.070 | 5.802 | 0.133 | -2.908 | 3.356 | 0.448 |
| chr3R:10638000-10644000 | 6000 | R-11 | CR44018 | R-11_CR44018 | 4.987 | 0.000 | 0.517 | 0.000 | 2.494 | 0.000 | nc | 2.318 | nc |
| chr2R:7229000-7235000 | 6000 | R116 | Or43a | R116_Or43a | 0.011 | 1.952 | 0.020 | 0.482 | 0.982 | 170.779 | 7.416 | -6.451 | 0.965 |
| chr3L:18505000-18511000 | 6000 | R117 | intergenic | R117_intergenic | 0.023 | 2.867 | 0.040 | 0.215 | 1.445 | 125.402 | 6.970 | -5.451 | 1.520 |
| chr3R:7815000-7821000 | 6000 | R118 | CG34384 | R118_CG34384 | 0.057 | 5.997 | 0.099 | 0.857 | 3.027 | 104.917 | 6.713 | -4.129 | 2.584 |
| chr2L:20833000-20839000 | 6000 | R119 | Spn38F | R119_Spn38F | 0.195 | 9.117 | 0.025 | 0.811 | 4.656 | 46.641 | 5.544 | -2.355 | 3.189 |
| chr2R:21175000-21181000 | 6000 | R120 | Pu | R120_Pu | 0.335 | 8.493 | 0.155 | 1.734 | 4.414 | 25.385 | 4.666 | -1.580 | 3.086 |
| chr3L:14272000-14278000 | 6000 | R121 | fz | R121_fz | 0.434 | 9.887 | 0.228 | 0.966 | 5.161 | 22.764 | 4.509 | -1.203 | 3.306 |
| chr3R:4531000-4537000 | 6000 | R122 | CG34357 | R122_CG34357 | 0.816 | 9.400 | 0.231 | 1.935 | 5.108 | 11.521 | 3.526 | -0.294 | 3.233 |
| chrX:22545000-22551000 | 6000 | R123 | CG17601 | R123_CG17601 | 0.872 | 7.050 | 0.048 | 0.529 | 3.961 | 8.087 | 3.016 | -0.198 | 2.818 |
