## Supplemental Table 5 for "The histone demethylase KDM3 prevents auto-immune piRNAs production in *Drosophila*"

| | MACS2 peaks | | Size (pb) | | % D.mel genome | | FC |
|---|---|---|---|---|---|---|---|
| **Enrichment (IP/INPUT)** | Control | Kdm3 GLKD | Control | Kdm3 GLKD | Control | Kdm3 GLKD | |
| H3K9me2 | 3983 | 9564 | 28029688 | 44185495 | 20.94 | 33.00 | 1.5764 |
| H3K9me3 | 2946 | 5282 | 23497658 | 27812726 | 17.55 | 20.77 | 1.1836 |
| Rhino | 467 | 1882 | 244106 | 1222674 | 0.18 | 0.91 | 5.0088 |

| Overlap | | | Size (pb) | | % D.mel genome | | |
|---|---|---|---|---|---|---|---|
| H3K9me2 | H3K9me3 | Rhino | Control | Kdm3 GLKD | Control | Kdm3 GLKD | |
| - | - | - | 105262523 | 84963645 | 78.62 | 63.46 | 0.8072 |
| + | + | - | 22694917 | 22676421 | 16.95 | 16.94 | 0.9992 |
| + | - | - | 5104765 | 20670476 | 3.81 | 15.44 | 4.0493 |
| - | + | - | 574297 | 4347392 | 0.43 | 3.03 | 7.5699 |
| + | + | + | 228444 | 788729 | 0.17 | 0.59 | 3.4526 |
| - | - | + | 14100 | 383892 | 0.01 | 0.29 | 27.2264 |
| + | - | + | 1562 | 49869 | 0.00 | 0.04 | 31.9264 |
| - | + | + | 0 | 184 | 0.00 | 0.00 | na |

**Extended Data Table 5. Genome wide ChIP-seq analyses using H3K9me2, H3K9me3 and Rhino antibodies.**
