## Supplemental Table 6 for "The histone demethylase KDM3 prevents auto-immune piRNAs production in *Drosophila*"

| | Size (pb) | | % new piRNA clusters sequences | | Fold change |
|---|---|---|---|---|---|
| | Control | Kdm3 GLKD | Control | Kdm3 GLKD | |
| H3K9me2 | 203141 | 1771831 | 8.53 | 74.38 | 8.7222 |
| H3K9me3 | 95781 | 747655 | 4.02 | 31.39 | 7.8059 |
| Rhino | 0 | 508852 | 0.00 | 21.36 | na |

| Overlap | | | Size (pb) | | % new piRNA clusters sequences | | Fold change |
|---|---|---|---|---|---|---|---|
| H3K9me2 | H3K9me3 | Rhino | Control | Kdm3 GLKD | Control | Kdm3 GLKD | |
| - | - | - | 2167497 | 373694 | 90.99 | 15.69 | 0.1724 |
| + | + | - | 84419 | 448252 | 3.54 | 18.82 | 5.3098 |
| + | - | - | 118722 | 1020055 | 4.98 | 42.82 | 8.5920 |
| - | + | - | 11362 | 31147 | 0.48 | 1.31 | 2.7413 |
| + | + | + | 0 | 268072 | 0.00 | 11.25 | na |
| - | - | + | 0 | 205144 | 0.00 | 8.61 | na |
| + | - | + | 0 | 35452 | 0.00 | 1.49 | na |
| - | + | + | 0 | 184 | 0.00 | 0.01 | na |

**Extended Data Table 6. New piRNA clusters ChIP-seq analyses using H3K9me2, H3K9me3 and Rhino antibodies.**
