## Supplemental Table 7 for "The histone demethylase KDM3 prevents auto-immune piRNAs production in *Drosophila*"

| | Size (kbp) | | Total size % | | 23-29nt unique mapper (rpm) | | | | | | Mean | | Std | | Density (RPKM) | | Density std | |
|---|---|---|---|---|---|---|---|---|---|---|---|---|---|---|---|---|---|---|
| | **Control** | **Kdm3 GLKD** | **Control** | **Kdm3GLKD** | **Control1** | **Control2** | **Control3** | **Kdm3 GLKD1** | **Kdm3 GLKD2** | **Kdm3 GLKD3** | **Control** | **Kdm3 GLKD** | **Control** | **Kdm3 GLKD** | **Control** | **Kdm3 GLKD** | **Control** | **Kdm3 GLKD** |
| **H3K9me2+H3K9me3+Rhi** | 0 | 268.072 | 0.00 | 11.25 | 0.000 | 0.000 | 0.000 | 1391.364 | 1430.378 | 1469.382 | 0.000 | 1430.375 | 0.000 | 39.009 | 0.00 | 5.34 | na | 0.146 |
| **Rhi (only)** | 0 | 205.144 | 0.00 | 8.61 | 0.000 | 0.000 | 0.000 | 687.522 | 724.654 | 756.968 | 0.000 | 723.048 | 0.000 | 34.751 | 0.00 | 3.52 | na | 0.169 |
| **H3K9me2+Rhi** | 0 | 35.452 | 0.00 | 1.49 | 0.000 | 0.000 | 0.000 | 123.389 | 129.435 | 122.002 | 0.000 | 124.942 | 0.000 | 3.952 | 0.00 | 3.52 | na | 0.111 |
| **H3K9me3 (only)** | 11.362 | 31.147 | 0.48 | 1.31 | 0.999 | 0.995 | 0.813 | 73.754 | 86.470 | 89.067 | 0.936 | 83.097 | 0.106 | 8.195 | 0.08 | 2.67 | 0.009 | 0.263 |
| **H3K9me2 (only)** | 118.722 | 1020.055 | 4.98 | 42.82 | 15.658 | 17.113 | 14.985 | 1710.886 | 1800.835 | 1846.301 | 15.919 | 1786.007 | 1.088 | 68.915 | 0.13 | 1.75 | 0.009 | 0.068 |
| **H3K9me2+H3K9me3** | 84.419 | 448.252 | 3.54 | 18.82 | 19.114 | 19.651 | 17.982 | 742.067 | 752.306 | 781.415 | 18.916 | 758.596 | 0.852 | 20.414 | 0.22 | 1.69 | 0.010 | 0.046 |
| **H3K9me3+Rhi** | 0 | 0.184 | 0.00 | 0.01 | 0.000 | 0.000 | 0.000 | 0.144 | 0.135 | 0.078 | 0.000 | 0.119 | 0.000 | 0.036 | 0.00 | 0.65 | na | 0.195 |
| **none** | 2167.497 | 373.694 | 90.99 | 15.69 | 109.231 | 136.047 | 108.919 | 46.602 | 42.187 | 22.531 | 118.066 | 37.107 | 15.573 | 12.815 | 0.05 | 0.10 | 0.007 | 0.034 |
| **Total** | 2382 | 2382 | | | 145.003 | 173.805 | 142.699 | 4775.729 | 4966.401 | 5087.743 | 153.836 | 4943.291 | 17.332 | 157.286 | 0.06 | 2.08 | 0.007 | 0.066 |

**Extended Data Table 7. piRNA production linked to the chromatin state of the 123 new piRNA clusters.**
